# The anti-sigma factor MucA is required for viability in *Pseudomonas aeruginosa*

**DOI:** 10.1101/2020.08.17.253617

**Authors:** Melissa C. Schofield, Daniela Rodriguez, Amanda A. Kidman, Erin K. Cassin, Lia A. Michaels, Elizabeth A. Campbell, Peter A. Jorth, Boo Shan Tseng

## Abstract

During decades-long infections in the cystic fibrosis (CF) airway, *Pseudomonas aeruginosa* undergoes selection. One bacterial genetic adaptation often observed in CF isolates is *mucA* mutations. MucA inhibits the sigma factor AlgU. Mutations in *mucA* lead to AlgU misregulation, resulting in a mucoid phenotype that is associated with poor CF disease outcomes. Due to its ability to be mutated, *mucA* is assumed to be dispensable for bacterial viability. Here we show that, paradoxically, a portion of *mucA* is essential in *P. aeruginosa*. We demonstrate that *mucA* is no longer required in a strain lacking *algU*, that *mucA* alleles encoding for proteins that do not bind to AlgU are insufficient for viability, and that *mucA* is no longer essential in mutant strains containing AlgU variants with reduced sigma factor activity. Furthermore, we found that overexpression of *algU* prevents cell growth in the absence of MucA, and that this phenotype can be rescued by overproduction of RpoD, the housekeeping sigma factor. Together, these results suggest that in the absence of MucA, the inability to regulate AlgU activity results in the loss of bacterial viability. Finally, we speculate that essentiality of anti-sigma factors that regulate envelope function may be a widespread phenomenon in bacteria.

## INTRODUCTION

The major cause of death in people with cystic fibrosis (CF), a human autosomal recessive genetic disease, is respiratory failure due to chronic lung infection. *Pseudomonas aeruginosa* is a prevalent CF respiratory pathogen (Cystic Fibrosis Foundation Patient Registry, 2019). The CF lung environment selects for mucoid *P. aeruginosa* mutants, which overproduce the exopolysaccharide alginate and are associated with poor disease prognosis (Douglas *et al*., 2009, Emerson *et al*., 2002, Farrell *et al*., 2009, Henry *et al*., 1992, Li *et al*., 2005, Nixon *et al*., 2001, Parad *et al*., 1999, Pedersen *et al*., 1992, Strateva *et al*., 2010). Conversion to the mucoid phenotype in clinical *P. aeruginosa* isolates, which is thought to be advantageous for chronic infection, is often caused by *mucA* mutations (Martin *et al*., 1993).

MucA is an anti-sigma factor to the alternative sigma factor AlgU (also known as AlgT, σ^E^, or σ^22^), which responds to envelope stress (Damron & Goldberg, 2012, Govan & Deretic, 1996). MucA is a transmembrane protein that sequesters AlgU away from RNA polymerase (RNAP) via its N-terminus (Li *et al*., 2019, Schurr *et al*., 1996, Xie *et al*., 1996). The C-terminus of MucA is in the periplasm, where it is protected from proteolysis by MucB (Mathee *et al*., 1997, Schurr *et al*., 1996). The envelope stress response is controlled via a regulated intramembrane proteolysis cascade: MucB dissociates from MucA upon stress detection, allowing the proteases AlgW and MucP to cleave MucA from the inner membrane (Qiu *et al*., 2007, Wood & Ohman, 2009). In the cytoplasm, MucA is further degraded by ClpXP, releasing AlgU to interact with RNAP and activate the AlgU regulon (Qiu *et al*., 2008). AlgU regulates at least 350 genes, including those responsible for the production of itself, MucA, and the alginate biosynthetic enzymes (Firoved & Deretic, 2003, Schulz *et al*., 2015, Wood & Ohman, 2009). This system is homologous to the well-studied envelope stress response in *Escherichia coli*: MucA shares 28% identity (72% similarity) to the anti-sigma factor RseA. The cognate sigma factor of the *E. coli* RseA, called RpoE, is homologous to and functionally interchangeable with AlgU (Yu *et al*., 1995), with 66% identity (93% similarity).

For several reasons, *mucA* is assumed to be dispensable for *P. aeruginosa* viability. First, after being released from the cell membrane, MucA is presumed to be fully degraded in the cytoplasm by ClpXP (Qiu *et al*., 2008). Second, *mucA* mutations often arise in CF clinical isolates (Boucher *et al*., 1997, Candido Cacador *et al*., 2018, Ciofu *et al*., 2008, Martin *et al*., 1993, Pulcrano *et al*., 2012), and many laboratory *mucA* mutants exist in the literature (Gallagher *et al*., 2011, Liberati *et al*., 2006, Skurnik *et al*., 2013, Turner *et al*., 2015). Third, there are three published strains in which the entirety of *mucA* is removed from the genome (Intile *et al*., 2014, Jones *et al*., 2010, Pritchett *et al*., 2015). Paradoxically, three different whole-genome studies in two different strains of *P. aeruginosa* identified *mucA* as an essential gene (Lee *et al*., 2015, Liberati *et al*., 2006, Skurnik *et al*., 2013). Additionally, it has been anecdotally noted that *mucA* could not be deleted from the *P. aeruginosa* PAO1 genome (Panmanee *et al*., 2019). To investigate this paradox, we systematically attempted to delete *mucA* from *P. aeruginosa* using allelic exchange. Our results show that a portion of *mucA* was required for bacterial viability in multiple *P. aeruginosa* strains, that *mucA* was no longer essential in a strain lacking *algU*, and that *mucA* alleles that encode for proteins that do not interact with AlgU were insufficient to rescue viability or led to a growth defect. We found that *algU* mutations encoding for a less active sigma factor could relieve *mucA* essentiality, and that AlgU overproduction in the absence of MucA was toxic. Interestingly, our works suggests that *mucA* essentiality can be suppressed by increasing the levels of the housekeeping sigma factor RpoD. Together, our results strongly suggest that the unregulated activity of AlgU itself, in the absence of MucA, leads to bacterial cell death.

## RESULTS

### *mucA* is essential for viability in a diverse set of *P. aeruginosa* isolates

To determine if *mucA* is essential, we attempted to delete the gene from various *P. aeruginosa* strains, using a modified allelic exchange protocol to turn it into a robust assay (Fig S1). An allele of *mucA* missing >95% of the coding region was introduced into *P. aeruginosa*. Using PCR-confirmed isolates containing both the endogenous and deletion alleles in the genome, we counter-selected for loss of one allele and then determined which *mucA* allele isolates resolved to via PCR. For non-essential genes, we should observe isolates that resolved to the endogenous or deletion allele. However, for essential genes, only cells that resolved to the endogenous allele should be isolated, as cells that resolved to the deletion allele should not survive in the absence of the gene. For statistical power, we screened ≥125 isolates for strains resolving only to the endogenous *mucA*. We performed this assay on a diverse set of wild-type *P. aeruginosa* strains, including four laboratory, five clinical and two environmental isolates (Table 1). These strains are not only isolated from diverse locations, but they also vary in their colony morphology and their exopolysaccharide production profile (Colvin *et al*., 2012). For all strains tested, all observed isolates resolved to the endogenous *mucA* allele (p < 0.0001, Fisher’s exact test), strongly suggesting that *mucA* is essential in wild-type *P. aeruginosa*. Since all tested strains required *mucA* for viability, we continued our experiments using PAO1 as a representative strain.

**Table 1.**
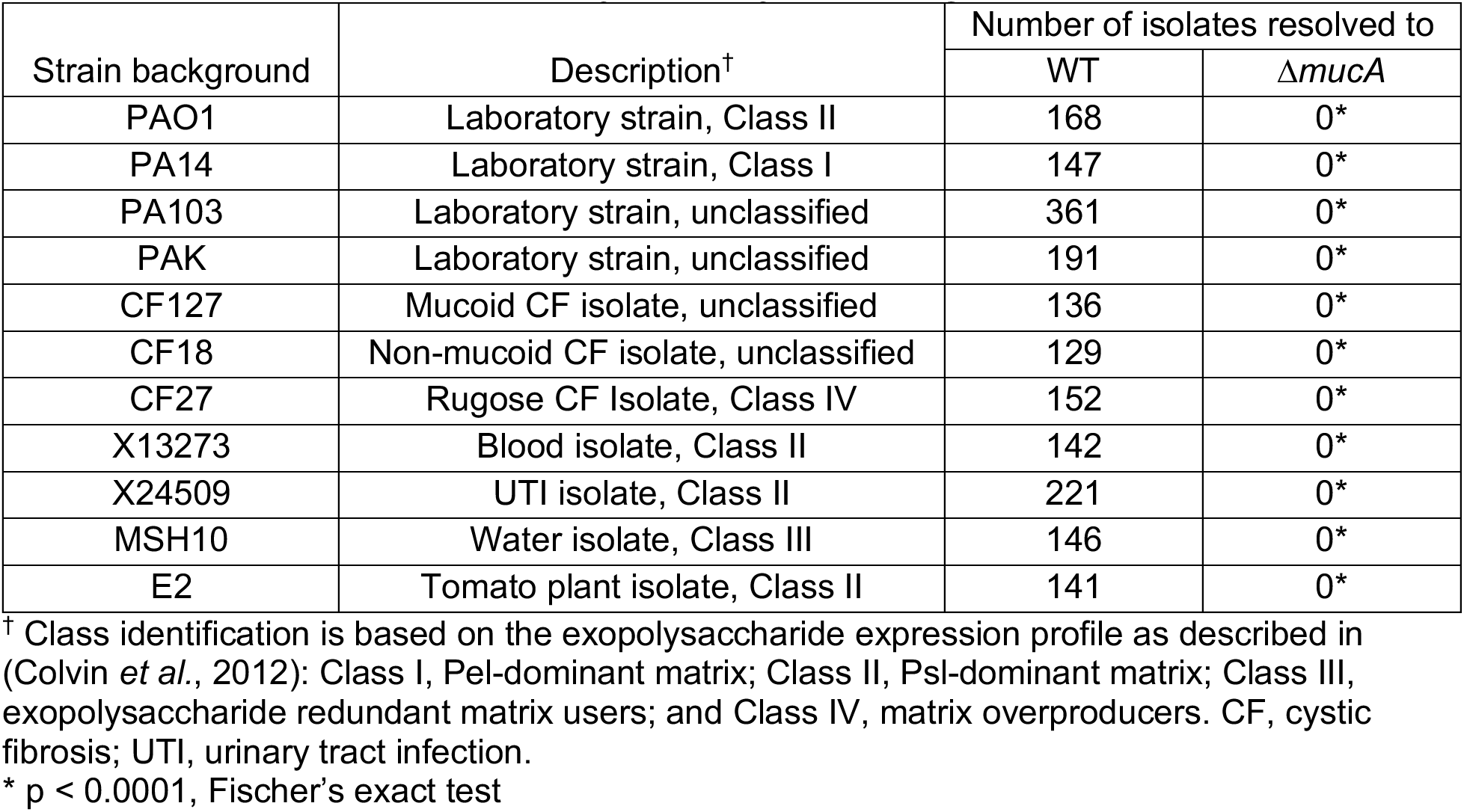
*mucA* is essential in a variety of wild-type *P. aeruginosa* strains.

### Depletion of MucA leads to cell death

To evaluate the effect of nutrient conditions on *mucA* essentiality, we tested the effect of depleting MucA on *P. aeruginosa* viability in different media. We engineered a strain lacking the native *mucA* and containing a chromosomally-integrated rhamnose-inducible copy (Δ*mucA att*Tn7::P*_rhaBAD_*-*mucA*). We determined viability of this strain grown without rhamnose over time. Since cells would cease producing MucA in the absence of rhamnose, the MucA present in the cells at the start of the experiment would be depleted as the cells divided. In all four media we tested, if rhamnose was not removed, the cells increased in density by ∼2-logs over time. In comparison, when rhamnose was removed, the cells lost viability over time in all four media (Fig 1), showing that *mucA* is essential in *P. aeruginosa* in various nutritional environments.

**Figure 1.**
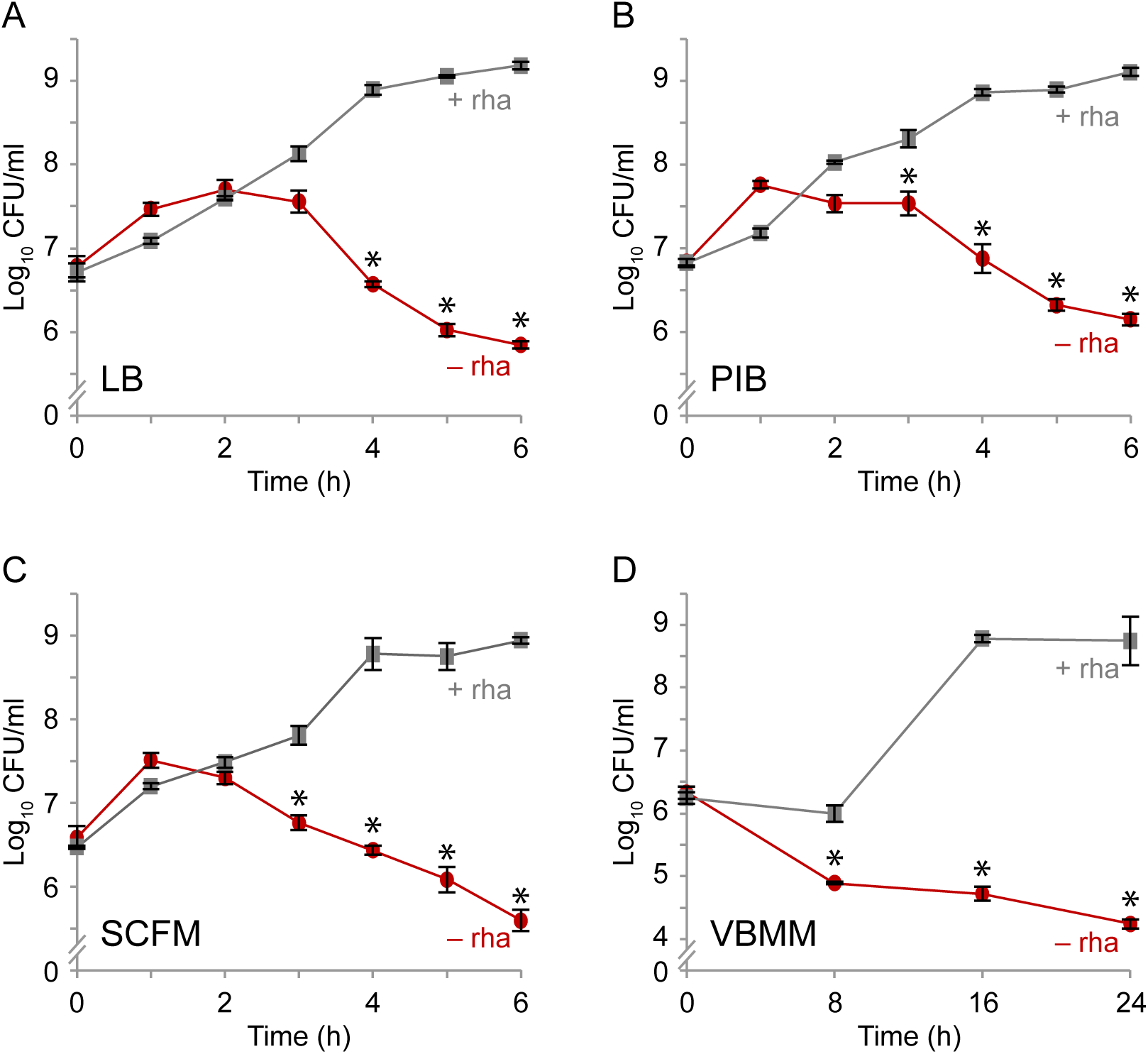
Depletion of MucA results in cell death. Viable colony counts of PAO1 Δ*mucA* attTn7::P*_rhaBAD_-mucA* over time in **(A)** LB, **(B)** PIB, **(C)** SCFM and **(D)** VBMM with (+ rha; gray squares) or without (– rha; red circles) 0.05% rhamnose. Viable colony counts after incubation in the indicated condition and time was determined by plating the cells on LB with 0.05% rhamnose to allow cells to recover and grow. Hash, broken y- axis; error bars, SEM (N=3). Asterisk, statistically different from that at the same time point grown in the presence of rhamnose (p < 0.01, N=3, mixed model ANOVA with post-hoc Bonferroni test).

### Alginate biosynthesis is not solely responsible for *mucA* essentiality

Clinical isolates with *mucA* mutations are mucoid due to the overproduction of alginate (Martin *et al*., 1993). To determine if alginate overproduction is responsible for *mucA* essentiality, we attempted to delete *mucA* from a strain lacking *algD*, a key alginate biosynthesis gene (Deretic *et al*., 1987). We were unable to delete *mucA* in this background (Table S1).

Expression of the alginate biosynthesis genes is controlled by three AlgU-regulated transcription factors that are active in mucoid cells: AlgB, AlgR, and AmrZ (Martin *et al*., 1994, Wozniak & Ohman, 1994, Wozniak *et al*., 2003). Since these transcription factors regulate many genes in addition to those involved in alginate biosynthesis (Huang *et al*., 2019, Jones *et al*., 2014, Kong *et al*., 2015, Leech *et al*., 2008), we tested if overexpression of these three regulons underlies *mucA* essentiality. We could not delete *mucA* from strains lacking these transcription factors (Table S1). We conclude that eliminating alginate biosynthesis and the expression of ∼50% AlgU regulon do not alleviate *mucA* essentiality.

### The first 50 amino acids of MucA are necessary and sufficient for cell viability

While we were unable to delete *mucA* using an allele lacking >95% of the coding region, *mucA* is often mutated in clinical isolates and many *mucA* transposon mutants exist (Fig S2), suggesting that only a portion of *mucA* is essential. Based on the co-crystal structures of MucA with AlgU and MucB (Li *et al*., 2019), the first 78 residues of MucA interact with AlgU, and the last 48 residues of MucA interact with MucB. We attempted to delete the endogenous *mucA* from a series of strains containing an ectopic chromosomally-integrated *mucA*, driven by its native promoter and encoding for truncation products (Fig 2A). We were unable to probe whether equivalent amounts of protein were produced across our strains, since our truncated versions of MucA are cleaved out of the membrane and degraded in the cytosol, as previously shown (Qiu *et al*., 2008). However, we expect that the ectopic alleles are expressed at similar levels prior to endogenous *mucA* deletion, as they are under the same promoter and at the same genomic site. These strains with two *mucA* alleles were non-mucoid, showing that the endogenous *mucA* allele, encoding full-length protein, is dominant over the ectopic alleles that encode for truncated MucA. We used a Δ*algD* background to make strains easier to manipulate, as deleting the endogenous *mucA* would result in alginate overproduction in some strains.

**Figure 2.**
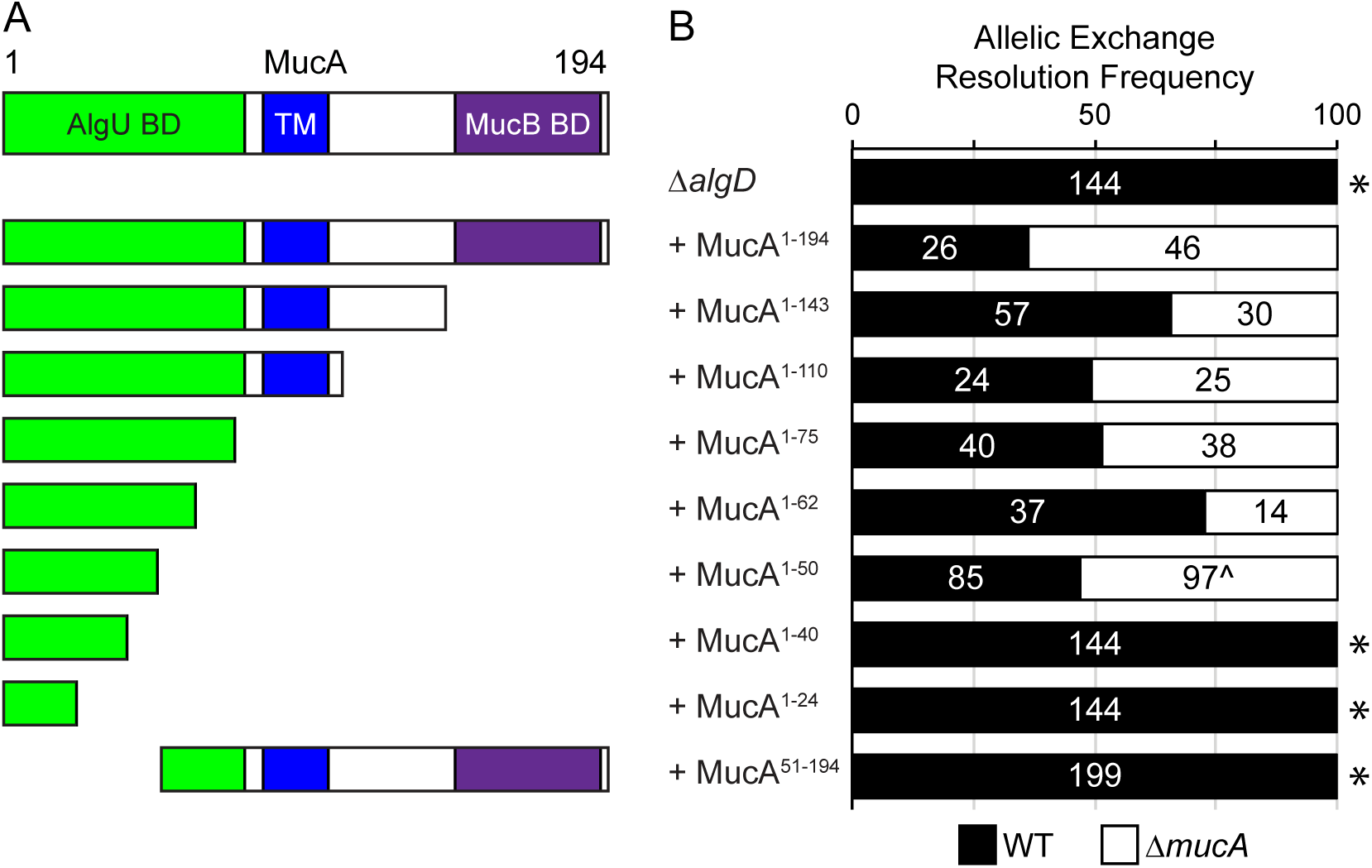
The first 50 amino acids of MucA are necessary and sufficient for viability. **(A)** Schematic of MucA encoded by the ectopic *mucA* alleles in PAO1 Δ*algD* strains tested for *mucA* essentiality in (B). MucA aa 1-143 lacks the MucB binding domain and is the product of the common *mucA22* allele. MucA 1-110 lacks the entire periplasmic domain. MucA aa 1-75 contains most of the AlgU binding domain, while shorter truncations contain only parts of the AlgU binding domain (MucA aa 1-62, 1-50, 1-40, and 1-24). Green, AlgU binding domain (AlgU BD); blue, the transmembrane domain (TM); and purple, the MucB binding domain (MucB BD). **(B)** Frequency of observed isolates resolving to the endogenous *mucA* allele (WT, black) or the deletion allele (Δ*mucA*, white) in the allelic exchange assay. Super-imposed on the bars are the number of isolates that were tested in each category. Asterisk, p < 0.0001; Fisher’s exact test. Caret, slower growing isolates.

We then attempted to delete the endogenous *mucA* from these strains. Since the flanking regions of the two alleles differ, our deletion allele specifically targets the native allele. We were able to easily recover isolates that resolved to the *mucA* deletion allele from strains containing ectopic alleles encoding for the full length MucA (aa 1-194), as well as MucA aa 1-143, aa 1- 110, aa 1-75, and aa 1-62 (Fig 2B). Interestingly, while the endogenous *mucA* could be deleted from strains containing an ectopic *mucA* that encoded for aa 1-50 (Fig 2B), isolates resolving to the deletion allele grew up more slowly than those resolving to the native allele. This suggests that while this allele is sufficient for viability, it is not well tolerated. All tested isolates of strains carrying *mucA* alleles encoding shorter products (aa 1-40 and aa 1-24) resolved to the native allele. Furthermore, we were unable to delete the native *mucA* from a strain with an ectopic *mucA* that encodes for aa 51-194 (Fig 2B). These results suggest that the first 50 amino acids of MucA are necessary and sufficient for *P. aeruginosa* cell viability. Consistent with these results, reported *mucA* mutations (Boucher *et al*., 1997, Candido Cacador *et al*., 2018, Ciofu *et al*., 2008, Martin *et al*., 1993, Pulcrano *et al*., 2012, Turner *et al*., 2015) almost entirely fall outside the region of *mucA* encoding the first 50 residues (Fig S2).

### The interaction of MucA with AlgU is required for cell survival

Since the only described function for MucA is to inhibit AlgU, we hypothesized that *mucA* essentiality is rooted in its regulation of AlgU. To test this, we attempted to delete *mucA* from a strain lacking *algU* (Δ*algU*). We were able to do so, with 16 of the tested isolates resolving to the deletion allele out of the 48 colonies tested (see Fig 4A), showing that *mucA* essentiality is *algU*- dependent.

Because our data suggest that the AlgU-binding domain of MucA is required for viability (Fig 2B), we hypothesized that the physical interaction between MucA and AlgU is necessary for cell survival. In the co-crystal structure (Li *et al*., 2019), four residues in MucA make more than one hydrogen bond with AlgU: D15, E22, R42, and E72 (Fig 3A). We engineered alleles that encode for MucA D15A, E22A, R42A, or E72A to maximally affect the hydrogen bonding while limiting the effects on overall protein structure. We used MucA aa 1-75 as a base because the co-crystal structure includes the first 78 residues of MucA. As described above, an allele encoding MucA aa 1-75 was sufficient for viability (Fig 2B). We were unable to delete the native *mucA* from strains carrying alleles encoding the D15A and R42A substitutions (Fig 3B). While we were able to delete the native *mucA* from the strain containing MucA E22A, isolates that resolved to the deletion allele grew up slower than those that resolved to the wild-type allele, suggesting that the cells do not tolerate this allele well. As expected, due to being outside the required region of MucA (Fig 2B), we were able to delete the native *mucA* from a strain containing MucA E72A with 9 of 48 tested isolates resolving to the deletion allele.

**Figure 3.**
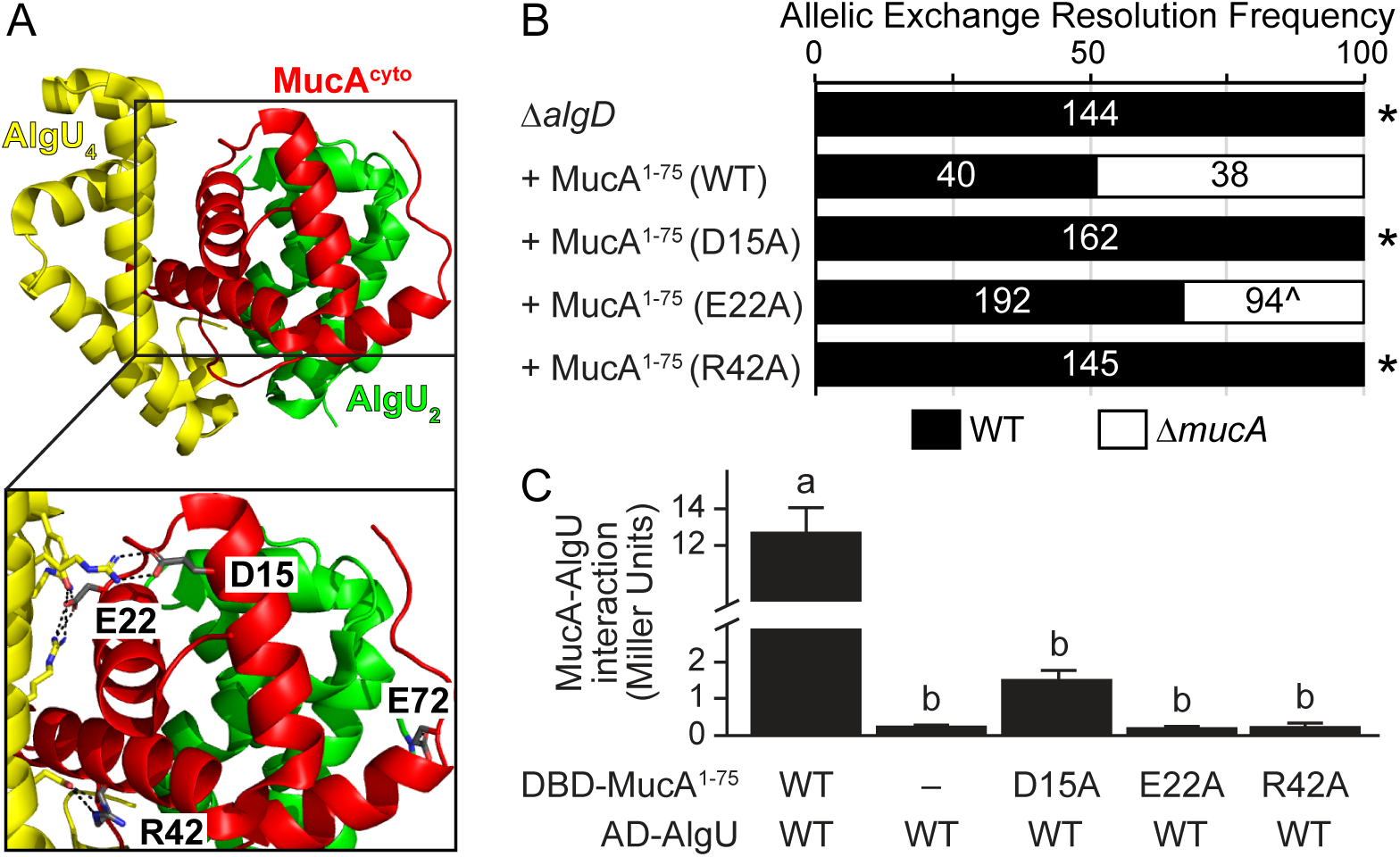
The physical interaction of MucA and AlgU is required for survival. **(A)** Four residues of MucA make greater than one predicted hydrogen bond with AlgU. The MucA-AlgU co-crystal structure (PDB 6IN7; Li et al., 2019) with the cytosolic domain of MucA (aa 1-78; red) and Regions 2 and 4 of AlgU (green, AlgU_2_; yellow, AlgU_4_) is shown. The residues of MucA that are predicted to make more than one hydrogen bond with AlgU (grey) are labeled in the inset. Black dotted lines, predicted hydrogen bonds; red atoms, oxygen; blue atoms, nitrogen. **(B)** Frequency of observed isolates resolving to the endogenous *mucA* allele (WT, black) or the deletion allele (Δ*mucA*, white) in the allelic exchange assay, using PAO1 Δ*algD att*Tn7::P*_algU_*-*mucA*, where *mucA* encodes for the indicated substitution. Super-imposed on the bars are the number of isolates that were tested in each category. Asterisk, p < 0.0001; Fisher’s exact test. Caret, slower growing isolates. **(C)** Substitution of MucA residues at its interface with AlgU abolish their binding via yeast two-hybrid. The first 75 residues of MucA were fused to the Gal4 DNA-binding domain (DBD-MucA^1-75^) and AlgU was fused to the Gal4 activation domain (AD-AlgU). Interaction of MucA and AlgU led to *lacZ* expression. Beta-galactosidase activity (in Miller units) was used as a proxy for the protein interaction strength. WT, wild-type protein sequence; –, no fusion protein included; hash, broken y-axis; error bars, SEM (N=3); letters, statistical groupings (p < 0.01; biological triplicate with technical quadruplicates; ANOVA with post-hoc Tukey HSD).

**Figure 4.**
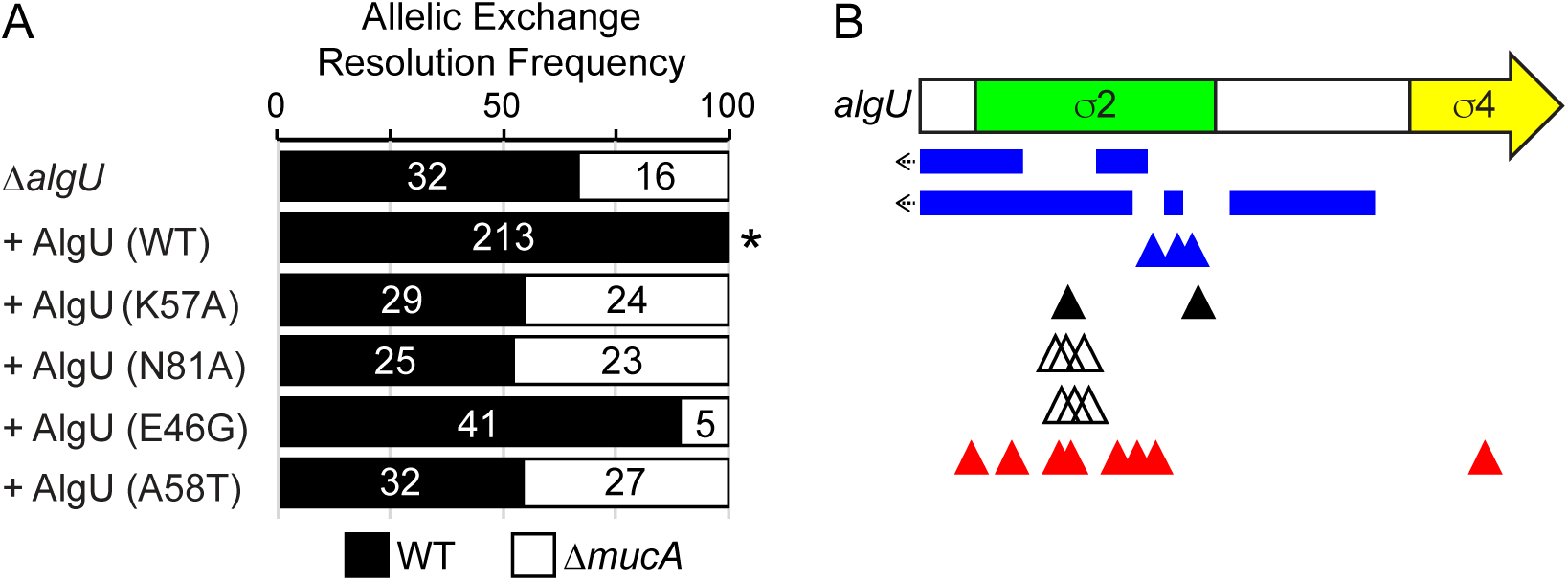
Mutations in AlgU can suppress *mucA* essentiality. **(A)** Frequency of observed isolates resolving to the endogenous *mucA* allele (WT, black) or the deletion allele (Δ*mucA*, white) in the allelic exchange assay, using PAO1 Δ*algU att*Tn7::P*_algU_*- *algU*, where *algU* encodes for the indicated substitution. Super-imposed on the bars are the number of isolates that were tested in each category. Asterisk, p < 0.0001; Fisher’s exact test. **(B)** Schematic of mutations seen in revertants that could grow in the absence of *mucA*. Revertants were selected by growing PAO1 Δ*mucA att*Tn7::P*_rhaBAD_*-*mucA* on media lacking rhamnose. Blue rectangles, multi-base pair deletions; left arrow, deletion extends into the promoter; blue triangles, single base pair deletions; black triangles, nonsense mutations; white triangles, duplications resulting in 3 or 4 amino acid insertions; red triangles, missense mutations. See Table S2 for a full description of the *algU* mutations.

To determine if the D15A, E22A, and R42A substitutions affect the ability of MucA to interact with AlgU, we used a yeast-two hybrid assay. Using beta-galactosidase activity as a proxy for their interaction, wild-type MucA aa 1-75 strongly interacted with AlgU (Fig 3C). This interaction was dependent on the presence of both MucA and AlgU, and the effect was not directional (Fig S3). In comparison, the D15A, E22A, and R42A MucA mutants failed to interact with AlgU (Fig 3C). We conclude that the interaction of MucA and AlgU are required for viability.

### Mutations that reduce AlgU activity alleviate the requirement for MucA

If *mucA* essentiality is due to its inhibition of the AlgU regulon, strains containing a mutant AlgU with lower affinity for DNA should alleviate *mucA* essentiality because the regulon expression is reduced in such mutants. We engineered Δ*algU* strains with an ectopic chromosomally- integrated *algU* allele encoding such DNA-binding mutants (K57A and N81A), based on homology to *E. coli* RpoE mutants that have decreased *in vitro* transcriptional activity (Campagne *et al*., 2014). As described above, we were able to delete *mucA* from a Δ*algU* strain (Fig 4A). This phenotype could be rescued via the ectopic addition of a wild-type *algU* allele, as we were no longer able to delete *mucA* from such a strain. In contrast, *mucA* could be deleted from strains carrying alleles encoding AlgU K57A or N81A, showing that these alleles failed to rescue the Δ*algU* phenotype (Fig 4A). To determine the effect of these substitutions on AlgU activity, we induced envelope stress using D-cycloserine (Wood *et al*., 2006) and measured AlgU activity using a plasmid-borne *gfp* reporter driven under the AlgU-regulated *algD* promoter (Damron *et al*., 2009). Similar to what is seen for RpoE (Campagne *et al*., 2014), these mutant strains had reduced AlgU activity upon induction of envelope stress (Fig 5A,B).

**Figure 5.**
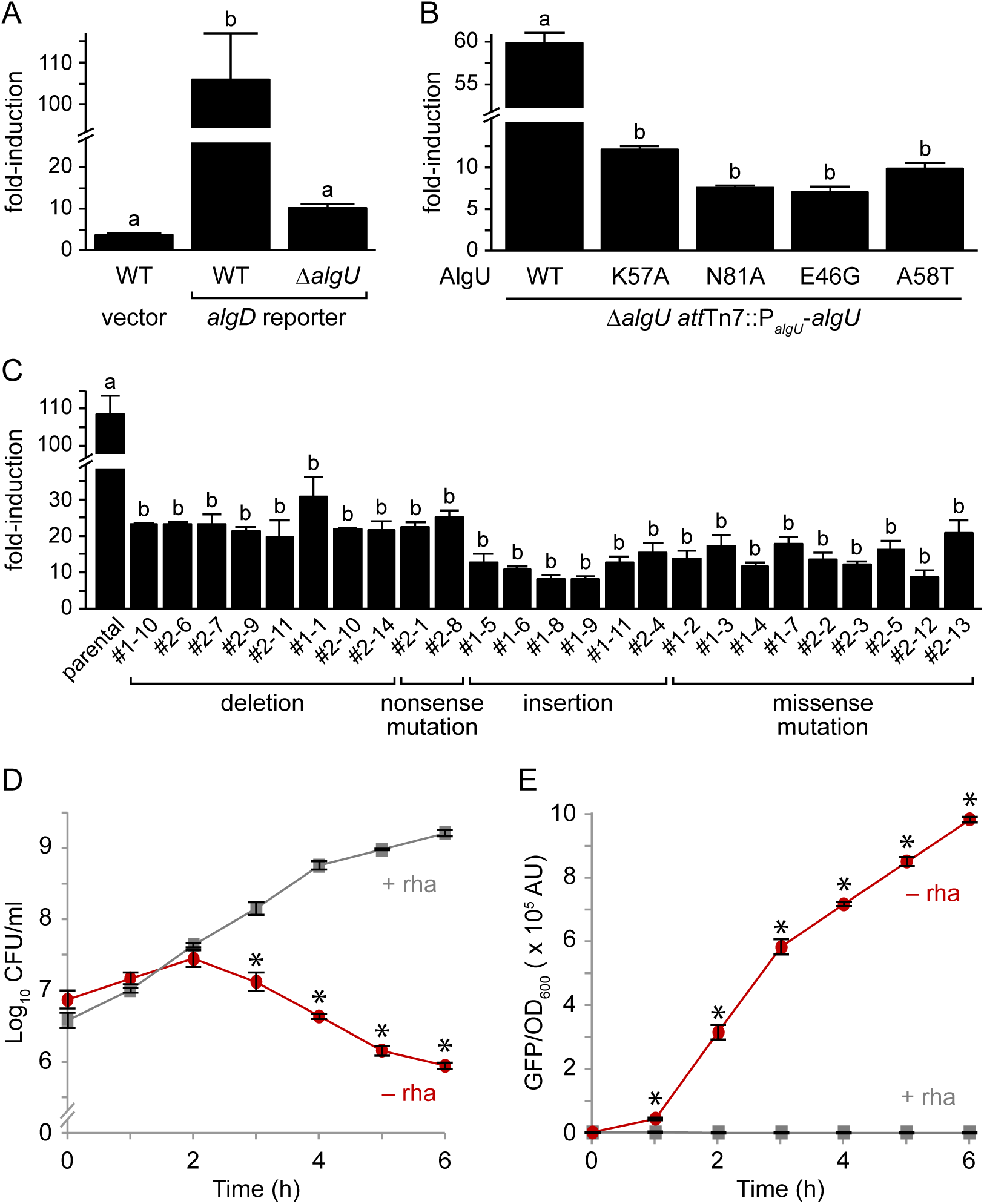
AlgU mutants that suppress *mucA* essentiality have decreased activity. **(A)** Fold-induction of GFP in PAO1 wild-type (WT) and Δ*algU* carrying a plasmid with either a promoter-less *gfp* (vector) or a *gfp* driven by the *algD* promoter (*algD* reporter), which is positively regulated by AlgU, after a 2h D-cycloserine treatment to activate envelope stress. GFP fluorescence was normalized to cell density and was then divided by the signal from untreated cells to determine the fold-induction. Hash, broken y-axis; error bars, SEM (N=3); letters, statistical groupings (p < 0.01; biological triplicate with technical quadruplicates; ANOVA with post-hoc Tukey HSD). **(B)** Fold-induction of GFP in PAO1 Δ*algU att*Tn7::P*_algU_*-*algU*, where *algU* encodes for the indicated substitution. The experiment and statistics are as described in (A). WT, the wild-type AlgU sequence. **(C)** Fold-induction of GFP in the revertants isolated from PAO1 Δ*mucA att*Tn7::P*_rhaBAD_*-*mucA* (parental) that can grow in the absence of MucA. The experiment and statistics are as described in (A), except all cells were grown in the presence of 0.05% rhamnose to allow for comparison to the parental strain, which grows only in the presence of rhamnose. The isolate identification numbers are shown and are grouped based on their *algU* mutation. **(D)** Viable colony counts of PAO1 Δ*mucA* attTn7::P*_rhaBAD_-mucA* carrying the *algD* reporter over time in LB with (+ rha; gray squares) or without (– rha; red circles) 0.05% rhamnose. Hash, broken y-axis; error bars, SEM (N=3). Asterisk, statistically different from that at the same time point grown in the presence of rhamnose (p < 0.01, N=3, mixed model ANOVA with post-hoc Bonferroni test). **(E)** Normalized GFP fluorescence of the samples in (D). GFP fluorescence was normalized to cell density. The statistical analysis is as described in (D).

Strains with the entirety of *mucA* deleted from *P. aeruginosa* PAO1, PAK, and PA103 exist (Intile *et al*., 2014, Jones *et al*., 2010, Pritchett *et al*., 2015). Since we were unable to delete *mucA* from these strain backgrounds (Table 1), we sequenced the genomes of these Δ*mucA* strains to determine if they contain suppressor mutations that allowed for their survival in the absence of *mucA*. In PAO1 Δ*mucA* (Pritchett *et al*., 2015), while the native *mucA* was deleted, the strain contained a second full-length copy of *algU* and a *mucA* allele that encoded for aa 1-155 elsewhere in the genome. As expected (Fig 2B), we were able to delete the native *mucA* allele from a PAO1 strain containing an ectopic allele that encoded for MucA aa 1-155 (with 5 of the 28 tested isolates resolving to the deletion allele), confirming that the ectopic *mucA* allele in the published PAO1 Δ*mucA* strain is sufficient for viability in the absence of the endogenous *mucA*. The PAK and PA103 Δ*mucA* strains (Intile *et al*., 2014, Jones *et al*., 2010) contain missense mutations in *algU,* which result in A58T and E46G, respectively. We replicated these mutations in PAO1 by inserting *algU* alleles encoding these substitutions in a Δ*algU* strain. We were able to delete *mucA* from these strains (Fig 4A). These results confirm that the *algU* alleles in the published PAK and PA103 Δ*mucA* strains suppress *mucA* essentiality. Using our reporter assay, we found that strains carrying these AlgU substitutions had reduced sigma factor activity (Fig 5B). We note that our reporter assay is not very sensitive to low levels of AlgU activity. Both PAK and PA103 Δ*mucA* strains are mucoid (Intile *et al*., 2014, Jones *et al*., 2010), suggesting that AlgU A58T and E46G are not completely inactive. Nevertheless, our results show that these AlgU mutants have significantly less transcriptional activity than the wild-type protein (Fig 5A,B).

To identify additional suppressors of *mucA* essentiality, we used the Δ*mucA att*Tn7::P*_rhaBAD_*- *mucA* strain. This strain was not viable in the absence of rhamnose, but natural revertants arose at a frequency of less than 1 in 10^9^ colony forming units. We sequenced the *algU* gene in 25 revertants, since AlgU mutants can suppress *mucA* essentiality (Fig 4A). All 25 isolates contained mutations in *algU* that are predicted to be hypomorphic (Fig 4B, Table S2). There were 10 revertants with deletions or nonsense mutations of *algU*, encoding either no product or a truncated product completely lacking Region 4 of the sigma factor. There were 6 revertants that contained multi-base pair duplications in *algU* that would lead to the insertion of 3 or 4 amino acids in Region 2 helix 3 of the sigma factor. There were 9 revertants containing missense mutations, 8 of which were unique, encoding the following substitutions: D18G, A21V, Y29C, A47T, D49G, Y59C, N81D, and R174G. Using a model of σ^E^ in complex with the RNAP core and the promoter element, we expect these insertions and substitutions to affect sigma factor folding, RNAP core interactions, or promoter interactions (Fig S4). Using our *algD* reporter assay in these natural revertants, we saw that all 25 revertants had much lower AlgU activity than the parental strain upon induction of envelope stress (Fig 5C), as predicted. Together, these data strongly suggest that *algU* mutations that encode for a protein with reduced transcriptional activity allow *P. aeruginosa* to survive in the absence of *mucA*.

If *mucA* essentiality is due to AlgU inhibition, AlgU activity should increase as MucA is depleted from cells, concomitant with the decrease in viability. To test this, the viability and fluorescence (as a proxy for AlgU activity) of a Δ*mucA att*Tn7::P*_rhaBAD_*-*mucA* strain containing our *algD* reporter plasmid was measured over time in the absence of rhamnose. This strain had a similar viability to that seen in Figure 2 (Fig 5D), and as MucA was depleted, the AlgU activity increased (Fig 5E). These results show the cell death observed upon MucA depletion is correlated with a dramatic increase in AlgU activity.

### Overexpression of *algU* in the absence of *mucA* leads to a growth defect

Our results suggest that *mucA* essentiality is due to unregulated AlgU activity. We reasoned that overexpression of *algU* should be toxic. However, *algU* can be overexpressed in wild-type *P. aeruginosa* that contain *mucA* (Qiu *et al*., 2008, Schulz *et al*., 2015). We therefore examined the effect of overproducing AlgU in wild type, Δ*algU*, and Δ*algU* Δ*mucA* strains, using a chromosomally integrated arabinose-inducible *algU*. Of note, the expression of *algU* is completely dependent on the inducer in the Δ*algU* and Δ*algU* Δ*mucA* strains, which lack positive feedback of AlgU on its own expression. Our results show that in the absence of inducer when AlgU is not overexpressed, all three strains had similar growth rates (Fig 6A). In the presence of 1% arabinose (i.e. high expression of *algU*), all three strains had a growth defect with the strain lacking *mucA* failing to grow at all.

**Figure 6.**
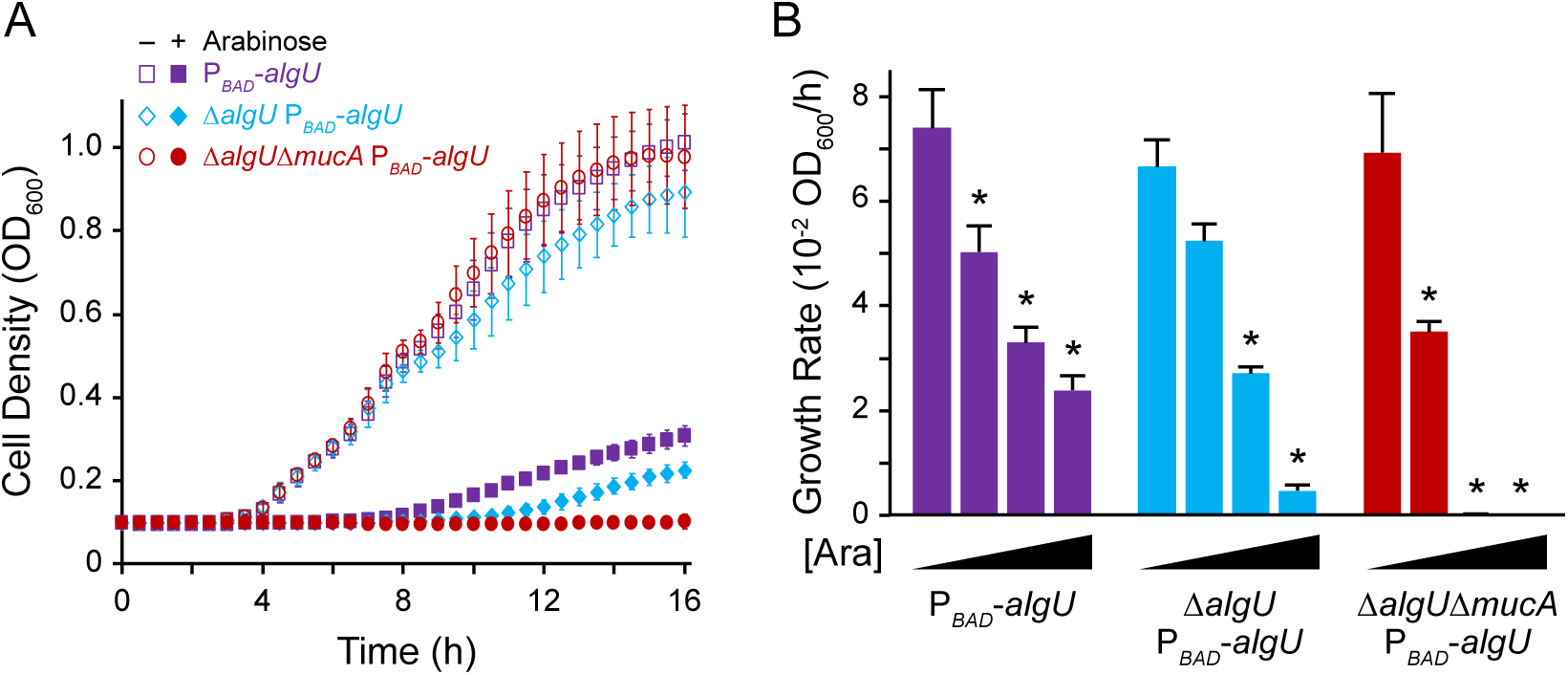
Overexpression of *algU* leads to a growth defect. **(A)** Growth curves of PAO1 strains containing an arabinose-inducible copy of *algU* in wild-type (purple squares), Δ*algU* (blue diamonds), and Δ*algU* Δ*mucA* (red circles) backgrounds in LB. Open symbols represent conditions in the absence of arabinose; closed symbols, with 1% arabinose. Error bars, SD (n=18). **(B)** Growth rate of strains as described in (A), with increasing induction of *algU*. Strains were grown in LB with 0%, 0.1%, 0.25%, and 1% arabinose ([Ara]). Error bars, SEM (N=3). Asterisk, statistically different from the same strain grown without arabinose (p < 0.01, N = 3, two-way ANOVA with post-hoc Bonferroni). See Table S3 for full statistical comparisons.

To determine if the Δ*algU* Δ*mucA att*Tn7::P*_araBAD_*-*algU* strain can grow under lower levels of *algU* induction, we tested a range of inducer concentrations (Fig 6B, Table S3). We found that although there was a growth defect, Δ*algU* Δ*mucA att*Tn7::P*_araBAD_*-*algU* was able to grow in the presence of 0.1% arabinose. The drop in growth rate in comparison to the no arabinose condition was statistically larger for the Δ*algU* Δ*mucA att*Tn7::P*_araBAD_*-*algU* than for the other two strains (N=3, p < 0.05, ANOVA with post hoc Tukey HSD), since this strain lacks the ability to produce any MucA to reduce AlgU activity. These results strongly suggest that in the absence of *mucA*, while a certain low level of AlgU activity is tolerated, high AlgU activity is fatal to the cell.

The above experiments were performed using only one medium. To determine if *algU* overexpression in the absence of *mucA* leads to a growth defect under other nutrient conditions, we determined the growth rate of the Δ*algU* Δ*mucA att*Tn7::P*_araBAD_*-*algU* strain in other media. The Δ*algU* Δ*mucA att*Tn7::P*_araBAD_*-*algU* strain failed to grow in the presence of 1% arabinose for all four media we tested (Fig S5), showing that AlgU overproduction is toxic under various nutrient conditions.

### Expression of *rpoD* can rescue the growth defect of *algU* overexpression

AlgU competes for RNAP binding with RpoD, the essential primary sigma factor (Yin *et al*., 2013). We, therefore, tested if the growth defect of high *algU* expression in the Δ*algU* Δ*mucA* background could be ameliorated by overexpressing *rpoD* under the same arabinose-inducible promoter. We tested growth in the presence of 2% arabinose, since these new strains contain two arabinose-inducible promoters, in contrast to the strains described in Figure 6. We confirmed that RpoD was overproduced in these new strains at similar levels upon arabinose induction (Fig S6). As expected (Fig 6), Δ*algU* Δ*mucA att*Tn7::P*_araBAD_*-*algU* failed to grow in the presence of 2% arabinose. The Δ*algU* Δ*mucA att*Tn7::P*_araBAD_*-*algU att*CTX::P*_araBAD_*-*rpoD* strain, however, was able to grow with arabinose. Furthermore, the growth rate of this strain with arabinose was indistinguishable from that of Δ*algU* Δ*mucA att*CTX::P*_araBAD_*-*rpoD* (Table S4).

There are two major explanations for how sigma factor competition may lead to toxicity during *algU* overexpression. First, the AlgU overproduction may reduce housekeeping gene expression due to the limited availability of RNAP to interact with RpoD. Second, *algU* overexpression results in high AlgU activity, which may lead to toxic expression of the AlgU regulon. We, therefore, constructed a strain containing an inducible *algU* encoding a K57A substitution in the Δ*algU* Δ*mucA* background. AlgU K57A exhibited reduced sigma factor activity (Fig 5A,B) and based on its location in the crystal structure (Campagne *et al*., 2014), we do not expect this substitution to affect RNAP affinity. In the presence of arabinose, the Δ*algU* Δ*mucA att*Tn7::P*_araBAD_*-*algU* K57A strain exhibited a growth rate greater than that of the strain overexpressing wild-type AlgU, but significantly less than that of the parental Δ*algU* Δ*mucA* strain (Fig 7, Table S4). RpoD overproduction in this background returned growth to a statistically similar level as that of a strain overexpressing RpoD alone (Table S4). Assuming that the K57A substitution does not affect AlgU affinity for RNAP, these results suggest that AlgU overexpression may lead to toxic expression of the AlgU regulon.

**Figure 7.**
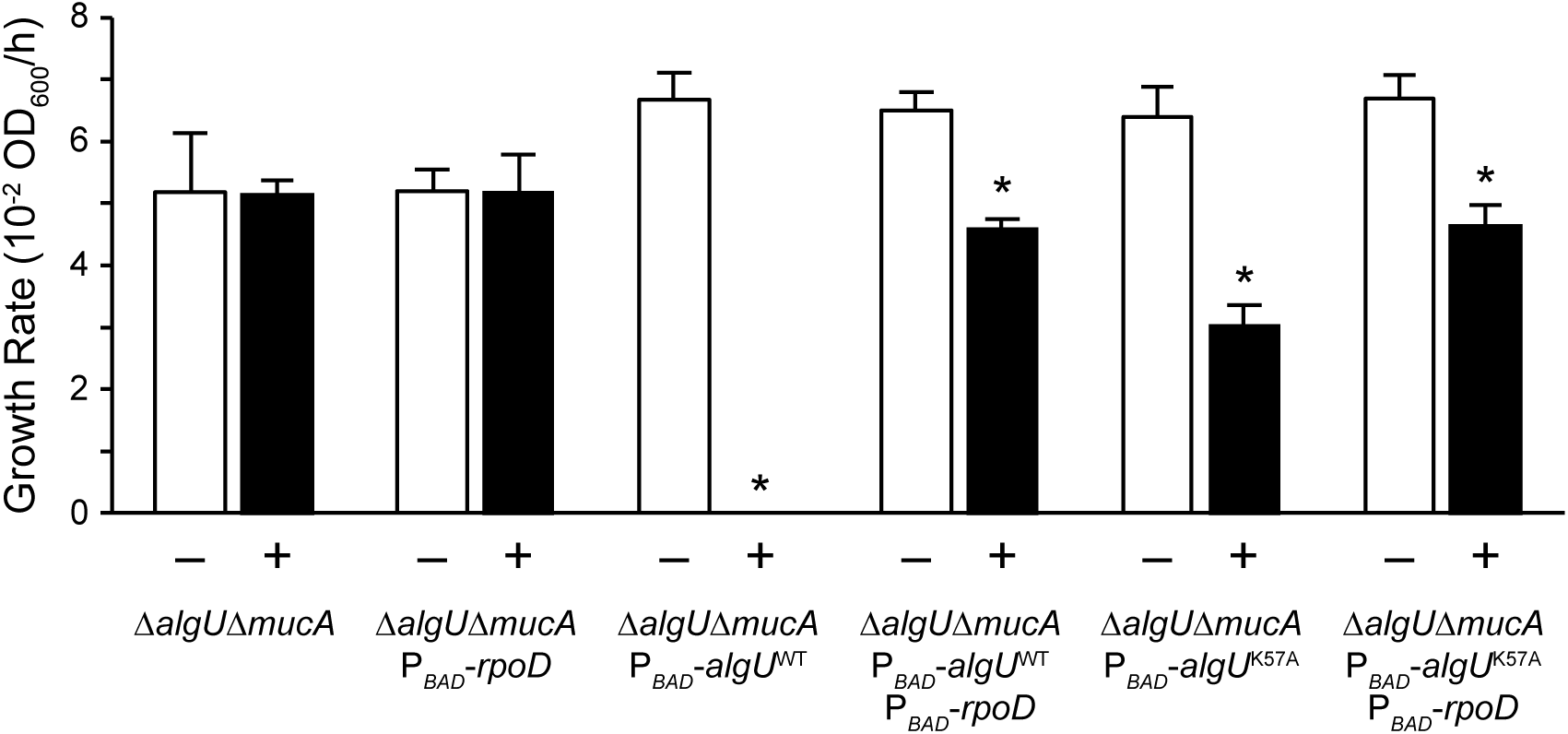
Growth defect caused by *algU* overexpression can be rescued via *rpoD* overexpression. Growth rate of indicated strains grown in LB with (+) or without (–) 2% arabinose. Error bars, SEM (N=3). Asterisk, statistically different from the same strain grown without arabinose (p < 0.01, N = 3, two-way ANOVA with post-hoc Bonferroni). See Table S4 for full statistical comparisons.

## DISCUSSION

Contrary to its assumed dispensability, our work shows that *mucA* is essential in a variety of *P. aeruginosa* wild-type strains and under various nutrient conditions (Table 1, Fig 1). Our data strongly suggest that unchecked AlgU activity in the absence of MucA leads to cell death (Fig 8). Under non-stress conditions, MucA inhibits AlgU. Under envelope stress or in strains containing *mucA* mutations, the anti-sigma factor is cleaved. We propose that while this cleaved cytosolic form of MucA does not inhibit AlgU to the same extent as the full-length protein, it is still able to inhibit AlgU to some degree. Although AlgU is active under such conditions, because *mucA* is positively regulated by AlgU, this negative feedback keeps AlgU activity under control. In comparison, this ability to control the positive feedback of AlgU on its own expression and activity is lost in the absence of MucA, which we propose is the cause of cell death. Supporting this model are several lines of evidence. First, *mucA* essentiality is rooted in its interaction with *algU* (Fig 2-3), strongly suggesting that the ability of MucA to inhibit AlgU is required for viability. Second, AlgU mutants with decreased transcriptional activity can suppress *mucA* essentiality (Fig 4-5), supporting the idea that decreasing the positive feedback of AlgU on its own expression allows for survival in the absence of MucA. Lastly, *algU* overexpression led to a growth defect, which was lethal at high levels in the absence of *mucA* (Fig 6), suggesting that high AlgU activity leads to cell death when MucA is not present. Overall, our work strongly suggests that *mucA* essentiality is caused by unchecked AlgU activity, in agreement with previous studies suggesting that overproduction of AlgU is toxic (Cross *et al*., 2020, Hershberger *et al*., 1995, Schurr *et al*., 1994).

**Figure 8.**
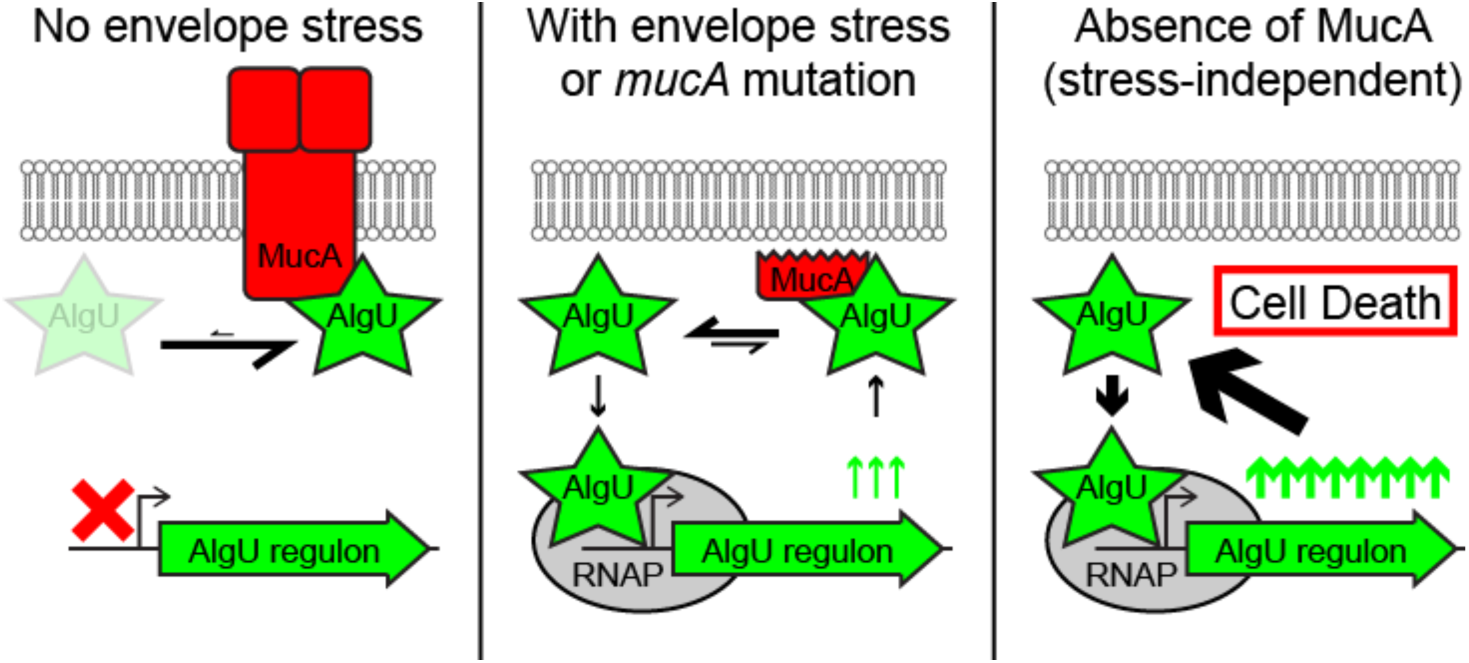
Unchecked AlgU activity in the absence of MucA leads to bacterial cell death. Under conditions of no envelope stress (left), the full-length MucA (red) binds strongly to AlgU (green star), leaving very little free AlgU to interact with the RNAP (grey oval). Therefore, the AlgU regulon (green arrow) is in the “off” state (red X). Under conditions of envelope stress or in strains containing *mucA* mutations that lead to a truncated product (middle), a cleaved cytosolic form of MucA is produced. This form can still interact with and inhibit AlgU. However, the strength of the interaction is weaker than with the full-length MucA, allowing for a pool of free AlgU that can then bind to and recruit RNAP to the promoters of its regulon. Under such conditions, the AlgU regulon, which includes *algU* and *mucA*, is activated and in the “on” state (green up arrows). However, because of the feedback on MucA, negative regulation of AlgU activity is still present. In the absence of *mucA* (right), there is no MucA to inhibit AlgU. All of the AlgU is free to interact with the core RNAP. This leads to high expression of the AlgU regulon and overproduction of AlgU itself. Under such strong positive feedback and in the absence of the negative regulator, the unchecked AlgU activity leads to cell death.

AlgU overproduction may be toxic because of reduced housekeeping gene expression (due to sigma factor competition with RpoD) and/or the increased expression of the AlgU regulon. Increasing RpoD levels alleviate the toxicity in both cases, by increasing the RNAP-RpoD complexes and reducing the RNAP-AlgU complexes. Our data does not definitively distinguish between these two mechanisms, which may not be mutually exclusive. Supporting a role for decreased housekeeping functions in AlgU toxicity, *mucA* was still essential in a strain lacking the three major AlgU-regulated transcription factors AlgB, AlgR, and AmrZ (Table S1), which regulate ∼50% of the AlgU regulon (Huang *et al*., 2019, Jones *et al*., 2014, Leech *et al*., 2008, Schulz *et al*., 2015). However, we cannot exclude that a combination of genes in the other half of the AlgU regulon may be responsible for *mucA* essentiality. Supporting the hypothesis that increased AlgU regulon expression causes toxicity, mutations in *algU*, which is AlgU-regulated, suppress *mucA* essentiality (Fig 4). While this autoregulation of AlgU is important for *mucA* essentiality under physiological conditions, it does not lead to toxicity per se, since overexpressing *algU* in a strain lacking the positive feedback led to a severe growth defect (Fig 6). Under such conditions, overproduction of AlgU K57A, which has reduced sigma factor activity (Fig 5B), caused a less severe growth defect than wild-type AlgU overproduction (Fig 7). Assuming that RNAP affinity is not affected by this AlgU substitution, this suggests that high AlgU activity and the ensuing regulon expression may be the mechanism underlying toxicity. Work in *Bacillus subtilis* has shown that the toxicity of unregulated SigM, an alternative sigma factor, is due specifically to membrane protein overproduction, which is alleviated by a gain-of-function mutation to the membrane insertase YidC1 (Zhao *et al*., 2019b). Since only ∼80 of the ∼200 AlgU-regulated predicted membrane proteins are regulated by AlgB, AlgR, and AmrZ (Huang *et al*., 2019, Jones *et al*., 2014, Leech *et al*., 2008, Schulz *et al*., 2015), it is possible that a gain-of-function YidC mutant may suppress *mucA* essentiality in *P. aeruginosa*. Such an experiment would help distinguish between the potential mechanisms underlying AlgU toxicity.

Anti-sigma factor essentiality is not unique to *P. aeruginosa*. In the defined transposon mutant library for *Vibrio cholerae*, an interruption in the *mucA* homolog *rseA* does not exist, suggesting that this anti-sigma factor may be essential (Cameron *et al*., 2008). Similarly, the *Pseudomonas syringae mucA* homolog is deemed essential based on transposon insertion analysis (Helmann *et al*., 2019). Furthermore, in *Mycobacterium tuberculosis*, the gene encoding the anti-sigma factor RslA is identified as essential (Griffin *et al*., 2011). Similar to our results (Fig 4), *rslA* can be deleted from *M. tuberculosis* lacking the gene encoding its cognate sigma factor SigL, which regulates genes involved in cell envelope processes (Dainese *et al*., 2006). Finally, there are two anti-sigma factors, YhdL and YxlC, in *B. subtilis* that are necessary for bacterial viability (Horsburgh & Moir, 1999, Koo *et al*., 2017, Mendez *et al*., 2012). The *yhdL* essentiality is notable as this anti-sigma factor negatively regulates SigM, an envelope stress response sigma factor (Horsburgh & Moir, 1999). Its essentiality could be suppressed by overproduction of SigA, the *B. subtilis* housekeeping sigma factor (Zhao *et al*., 2019a) and by a hyperactive form of the membrane insertase YidC1 (Zhao *et al*., 2019b). Interestingly, the YidC1 mutant ameliorated the toxicity of unregulated SigM by reducing the secretion stress associated with high membrane protein production (Zhao *et al*., 2019b). Taken together with our data, while not universal (as there are no essential anti-sigma factors in *E. coli* (Baba *et al*., 2006)), we speculate that essentiality of anti-sigma factors that regulate cell envelope function may be a widespread phenomenon in bacteria and that their essentiality is due to the increased production of membrane proteins when their sigma factors are not held in check.

In clinical CF isolates of *P. aeruginosa*, *mucA* mutations that lead to C-terminal truncations are common (Fig S2) and are expected to result in proteins that retain partial AlgU-inhibitory function. Interestingly, there is one *P. aeruginosa* CF isolate that is reported to have a *mucA* mutation that would result in a truncation within the first 50 aa of the protein (Boucher *et al*., 1997). Our sequencing data of the published Δ*mucA* strains (Intile *et al*., 2014, Jones *et al*., 2010, Pritchett *et al*., 2015) suggests that this isolate may contain a suppressor mutation that allows it to survive in the absence of a functional MucA. Nonetheless, in CF clinical isolates, *mucA* mutations that lead to C-terminal truncations are the norm, while null mutations in *mucA* with a presumed hypomorphic *algU* allele is rare (Fig S2). Furthermore, our results (Fig 4) agree with the literature showing that mucoid isolates with *mucA* mutations revert to a non-mucoid state via changes to *algU* (Ciofu *et al*., 2008, DeVries & Ohman, 1994, Sautter *et al*., 2012, Schurr *et al*., 1994). While these *algU* secondary site mutations are found in non-mucoid CF isolates with *mucA* mutations, such isolates are detected at a lower frequency (Ciofu *et al*., 2008). As suggested by Ciofu and associates, this may be due to the importance of AlgU for the survival of *P. aeruginosa* in the CF lung environment, suggesting that reducing AlgU activity may increase *P. aeruginosa* eradication from the CF airway. We propose that the MucA-AlgU interaction may serve as a good therapeutic target. Our results show that destabilizing the MucA-AlgU interaction results in bacterial cell death or mutations in *algU* that result in reduced sigma factor activity and likely reduced mucoid conversion, of which either outcome could be beneficial in the treatment of *P. aeruginosa* CF lung infections.

## METHODS AND MATERIALS

### Bacterial strains and growth conditions

The bacterial strains, plasmids, and oligonucleotides used for this study are in Tables S5-S7. The construction of strains is described in Supplementary Information. Bacteria were grown at 37°C in LB with shaking or on semi-solid LB media, unless otherwise noted.

### Allelic exchange assay

The protocol, depicted in Figure S1, was modified from (Hmelo *et al*., 2015). Briefly, a vector containing the *mucA* deletion allele flanked by approximately equal length homology regions is introduced via conjugation. Merodiploids were selected on semi-solid VBMM (Vogel & Bonner, 1956) with 60 mg/L gentamicin. Using OMS118 and OMS119, PCR was performed on at least six isolates to confirm that the presence of both the wild-type and deletion alleles of *mucA*. Confirmed merodiploids were then individually streaked on NSLB (10 g/L tryptone, 5 g/L yeast extract) with 10% sucrose semi-solid media for counterselection. PCR was performed on eight colonies per merodiploid, using OBT601 and OBT602 to determine the resolution to either the endogenous or deletion allele.

### MucA depletion assay

Cells were grown in LB with 0.05% rhamnose at 37°C with shaking to an OD_600_ of 0.3. After washing, the culture was then divided in two, half resuspended with rhamnose and half without in its respective media base (LB, PIB, SCFM, or VBMM). Cells were incubated at 37°C with shaking. At the indicated time points, two aliquots were removed from each culture, serially diluted and plated in triplicate onto LB agar plates with 0.05% rhamnose for recovery. Colonies were counted and log_10_-transformed CFU/mL of the culture was calculated.

### Yeast two-hybrid assay

The ProQuest Two-Hybrid System (Invitrogen) was used, per manufacturer’s instructions. Briefly, yeast containing the Gal4 activation domain-based prey and the Gal4 DNA-binding domain-based bait vectors were grown overnight in SD-Leu-Trp broth (Clontech). The OD_600_ value was recorded. ONPG (VWR) was added to lysed cells, and the mixture was incubated at 37°C until a light yellow color was achieved. The incubation time was recorded. The OD_420_ of the supernatants was determined using a Synergy Hybrid HTX Microplate Reader (Bio-Tek Instruments). Beta-galactosidase activity was determined using Miller units, based on the following equation: 1000 x OD_420_/(time x culture volume x OD_600_).

### AlgU activity assay

Strains of interest were transformed with a plasmid-borne *algD* reporter (pBT435) via electroporation (Choi *et al*., 2006). Strains were grown overnight in LSLB with 50 mg/L gentamicin and 0.05% rhamnose (where indicated) at 37°C with shaking. The overnight culture was diluted 1:100 and grown to an OD_600_ of 0.3. Cells were then treated with fresh 400 mg/L D-cycloserine for 2 h at 37°C with shaking, as previously described (Wood *et al*., 2006). Cells were then pelleted and resuspended in PBS. GFP fluorescence and OD_600_ was determined using a Synergy Hybrid HTX Microplate Reader (Bio-Tek Instruments). To normalize the data, the GFP fluorescence was divided by the OD_600_ for each data point. To determine the fold induction, the ratios for the treated samples were divided by the average of that for the same strain not treated with D-cycloserine. The resulting ratios of the triplicate samples were averaged.

### Natural revertant assay

*P. aeruginosa* Δ*mucA att*Tn7::P*_rhaBAD_*-*mucA* was grown in LB with 0.05% rhamnose at 37°C with shaking to an OD_600_ of 1. Cells were washed and plated on semi-solid LB plates without rhamnose. Plates were incubated for 24-48 h at 37°C. For isolates that grew on LB without rhamnose, *algU* was Sanger sequenced.

## ACKNOWLEGDEMENTS

The authors would like to thank Cai Tao and Maria Alexandra Ledesma for technical assistance; and Joe J. Harrison for critical discussions of this work. MCS, AAK, DR, and BT are supported by the Cystic Fibrosis Foundation (TSENG19I0). MCS and EKC are supported by University of Nevada Las Vegas doctoral graduate research assistantships. EAC and PAJ are supported by the National Institutes of Health (R01GM114450 and K22AI127473, respectively). PAJ is also supported by the Cystic Fibrosis Foundation (JORTH17F5). The authors have no conflicts of interest to disclose.

## AUTHOR CONTRIBUTIONS

Conceptualization: MCS, BST; Acquisition, analysis, or interpretation of data: MCS, DR, AAK, EKC, LAM, EAC, PAJ, BST; Writing – original draft: MCS, BST; and Writing – review and editing: MCS, DR, LAM, EAC, PAJ, BST.

---

**Figure S1.**
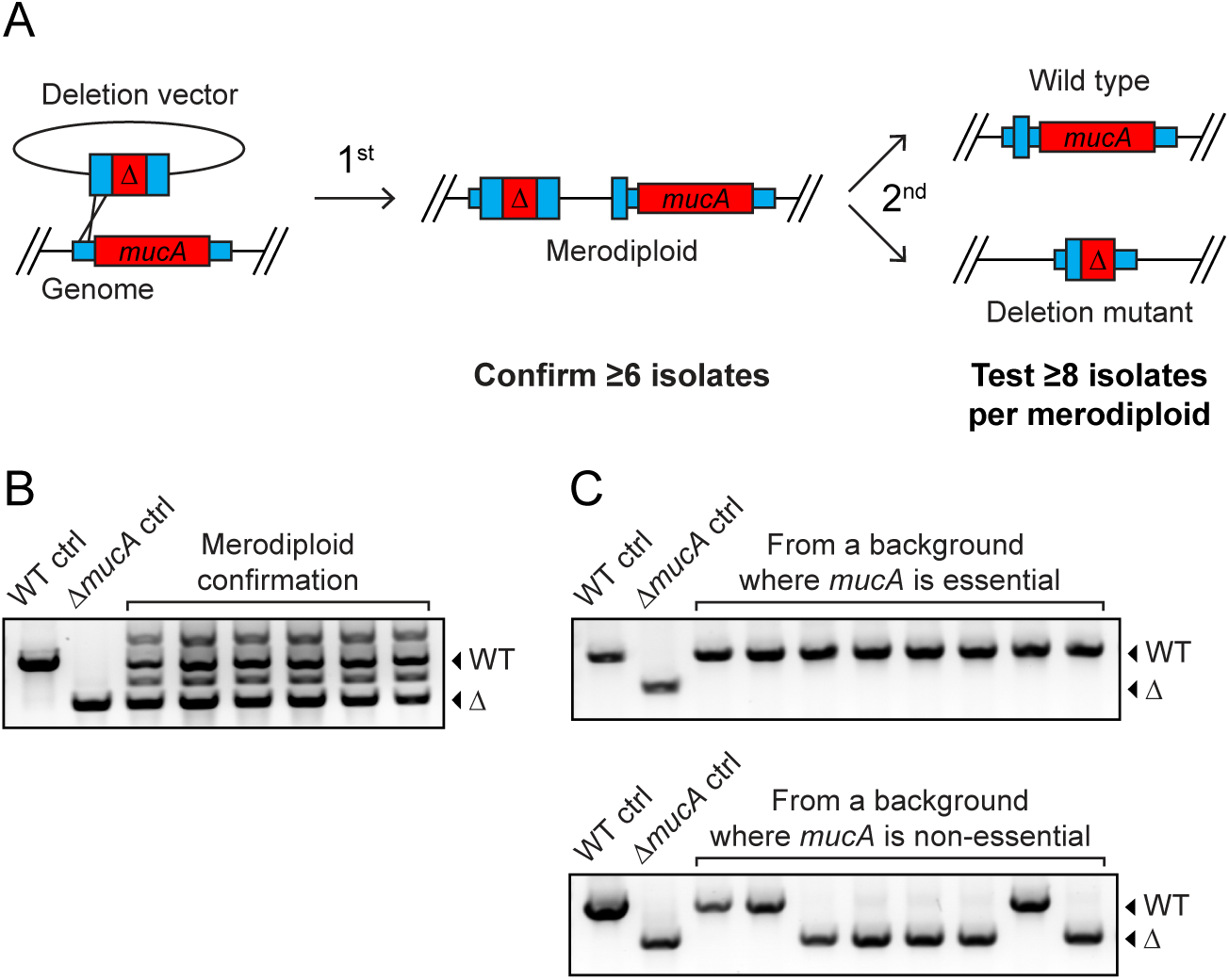
Allelic exchange assay. **(A)** Schematic of assay. To delete the endogenous *mucA* (short red box with *mucA*) from the genome, the deletion vector pBT396, which contains an allele of *mucA* that is missing >95% of the coding region (tall red box with delta), was introduced into *P. aeruginosa* via conjugation. Regions of homology (tall blue boxes) to the genome (short blue boxes) approximately 400 base pairs in length flank the deletion allele and allow for recombination into the genome (X) to create a merodiploid. This first recombination event (1^st^) can be selected for using antibiotics due to integration of the vector backbone, which contains a gentamicin resistance marker. At least six merodiploids were confirmed via PCR to ensure the genome contained both the deletion and endogenous *mucA* allele. Merodiploids can then undergo a second recombination event (2^nd^) that will lead to the loss of one of the two alleles, resolving to either the endogenous or deletion allele, which results in a cell that is either wild-type (top) or a deletion mutant (bottom), respectively. For each of the six confirmed merodiploids, we counter-selected for the second recombination event via the loss of *sacB*, a vector backbone marker. Using PCR, eight isolates per merodiploid were tested to determine which allele each isolate resolved to. If a gene is non-essential, we expect to observe both wild type or deletion mutants. However, if a gene is essential, we expect to isolate only those cells that resolved to wild type, as the cells cannot survive with the deletion allele. If we are unable to delete *mucA* from the first replicate of this experiment, we perform two additional biological replicates, for a minimum total of 125 colonies screened. If all isolates resolve to wild-type, we deem *mucA* to be essential in that strain background (p < 0.0001, Fisher’s exact test). **(B)** Representative image of merodiploid confirmation. The PCR products corresponding to the endogenous allele and the deletion allele are indicated with an arrow labelled “WT” and “Δ,” respectively. Positive (WT ctrl; PAO1) and negative (Δ*mucA* ctrl; BTPa355) controls are included. PCR products from six representative merodiploids are shown. **(C)** Representative image of PCR products from isolates after the second recombination. Image is labeled as in (B). Top, the PCR products of 8 isolates from a strain in which *mucA* was deemed essential. Bottom, the PCR products of 8 isolates from a strain in which *mucA* was not essential.

**Figure S2.**
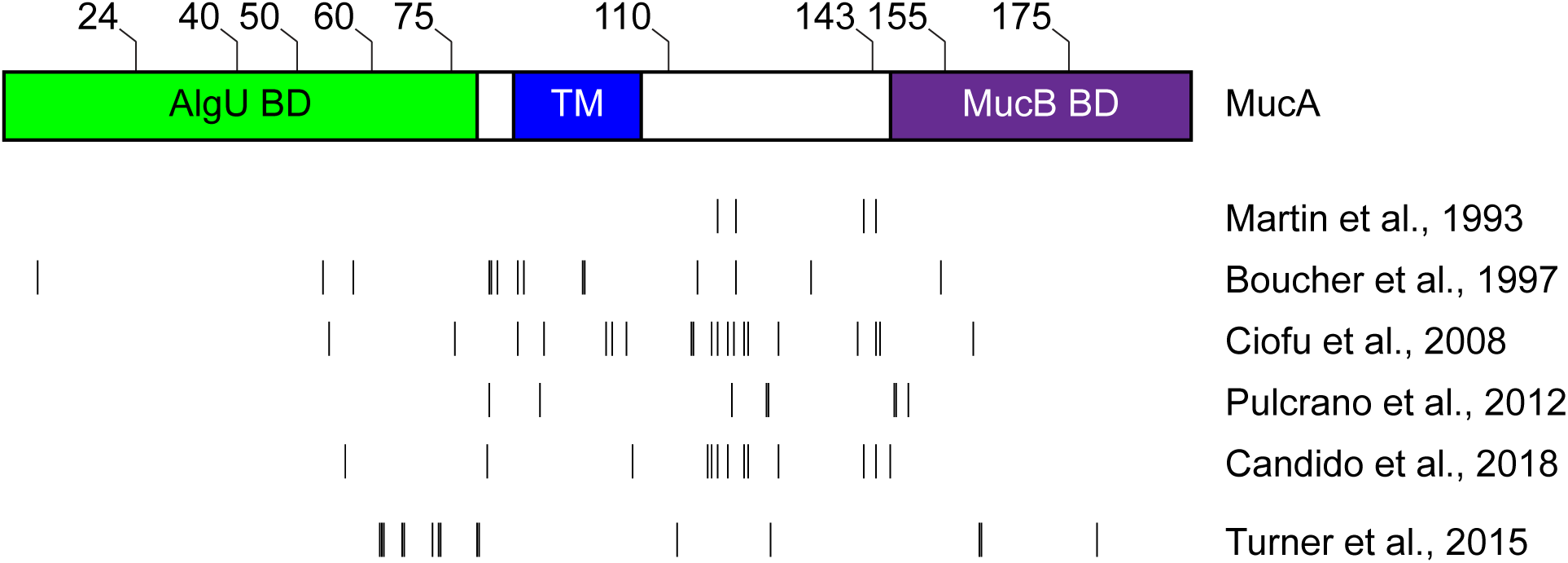
Published mutations in *mucA*. Top, schematic of MucA with the regions indicated encoding for the AlgU binding domain (green, AlgU BD), the transmembrane domain (blue, TM), and the MucB binding domain (purple, MucB BD). Hash marks on top indicate the residue number in MucA. Bottom, sites of *mucA* mutations that lead to the production of a truncated protein in published isolates. Each line represents one or more individual isolates described in the references on the right. The mutations for the upper five references (Boucher *et al*., 1997, Candido Cacador *et al*., 2018, Ciofu *et al*., 2008, Martin *et al*., 1993, Pulcrano *et al*., 2012) are in cystic fibrosis clinical isolates of *P. aeruginosa*, while the mutations in the last reference (Turner *et al*., 2015) are random transposon insertions in the laboratory PAO1 and PA14 strains.

**Figure S3.**
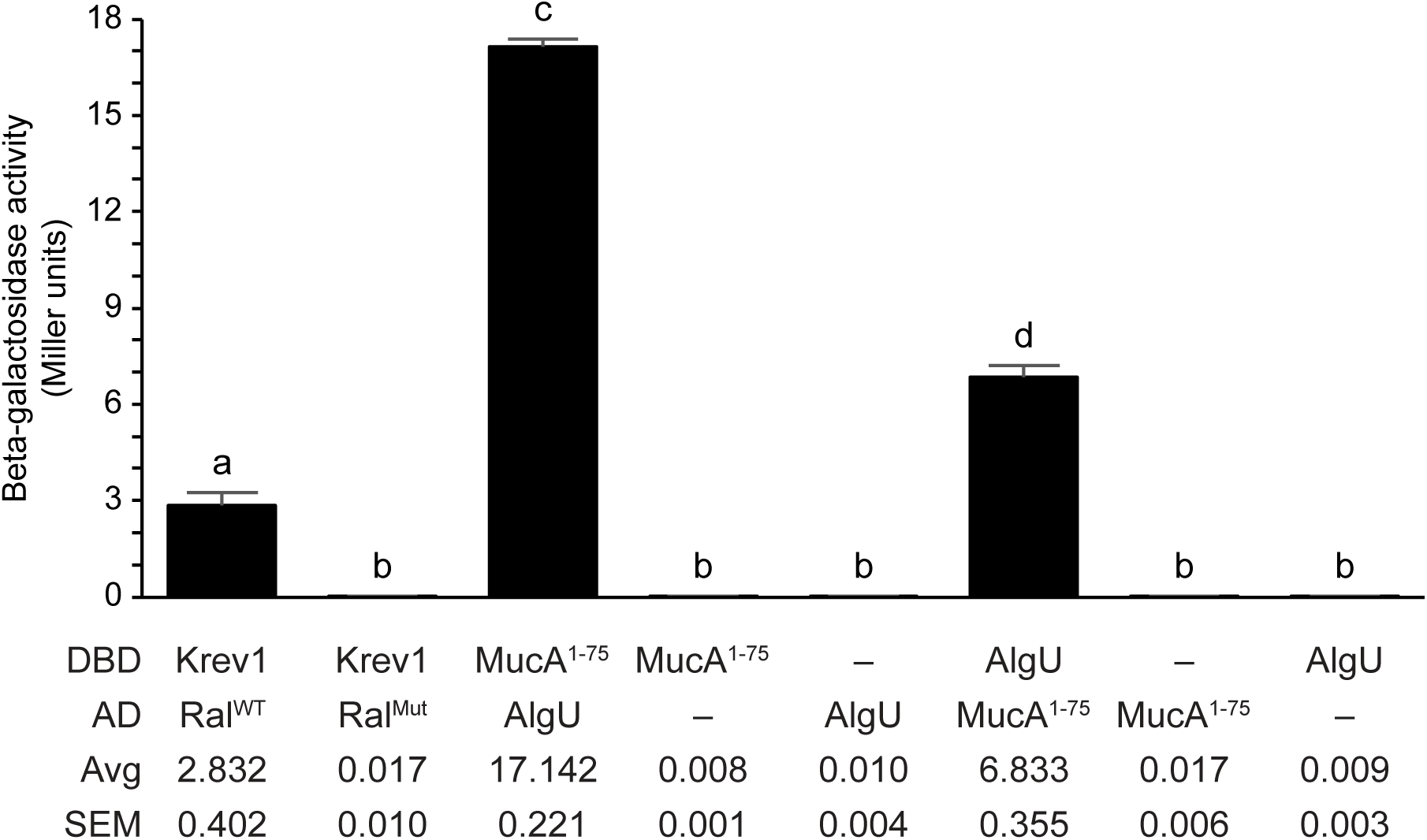
MucA aa 1-75 and AlgU interact via yeast two-hybrid assay. The indicated proteins were fused to the Gal4 DNA binding domain (DBD; “bait”) or the Gal4 activation domain (AD; “prey”). Interaction of the bait and prey proteins drive the expression of *lacZ*. Beta-galactosidase activity (in Miller units) was used as a proxy for the protein interaction strength. The average (Avg) and standard error of the mean (SEM) of each bar are indicated (N=3). As positive and negative controls, Krev1 is known to interact with wild-type RalGDS (Ral^WT^), but not the mutant RalGDS (Ral^Mut^) via yeast two-hybrid. AlgU, construct encoding full- length wild-type AlgU; MucA^1-75^, construct encoding only the first 75 residues of wild-type MucA; –, no fusion protein included; error bars, SEM (N=3); letters, statistical groups with the different letters representing statistically different groups (p < 0.01; biological triplicate with technical quadruplicates; ANOVA with post-hoc Tukey HSD).

**Figure S4.**
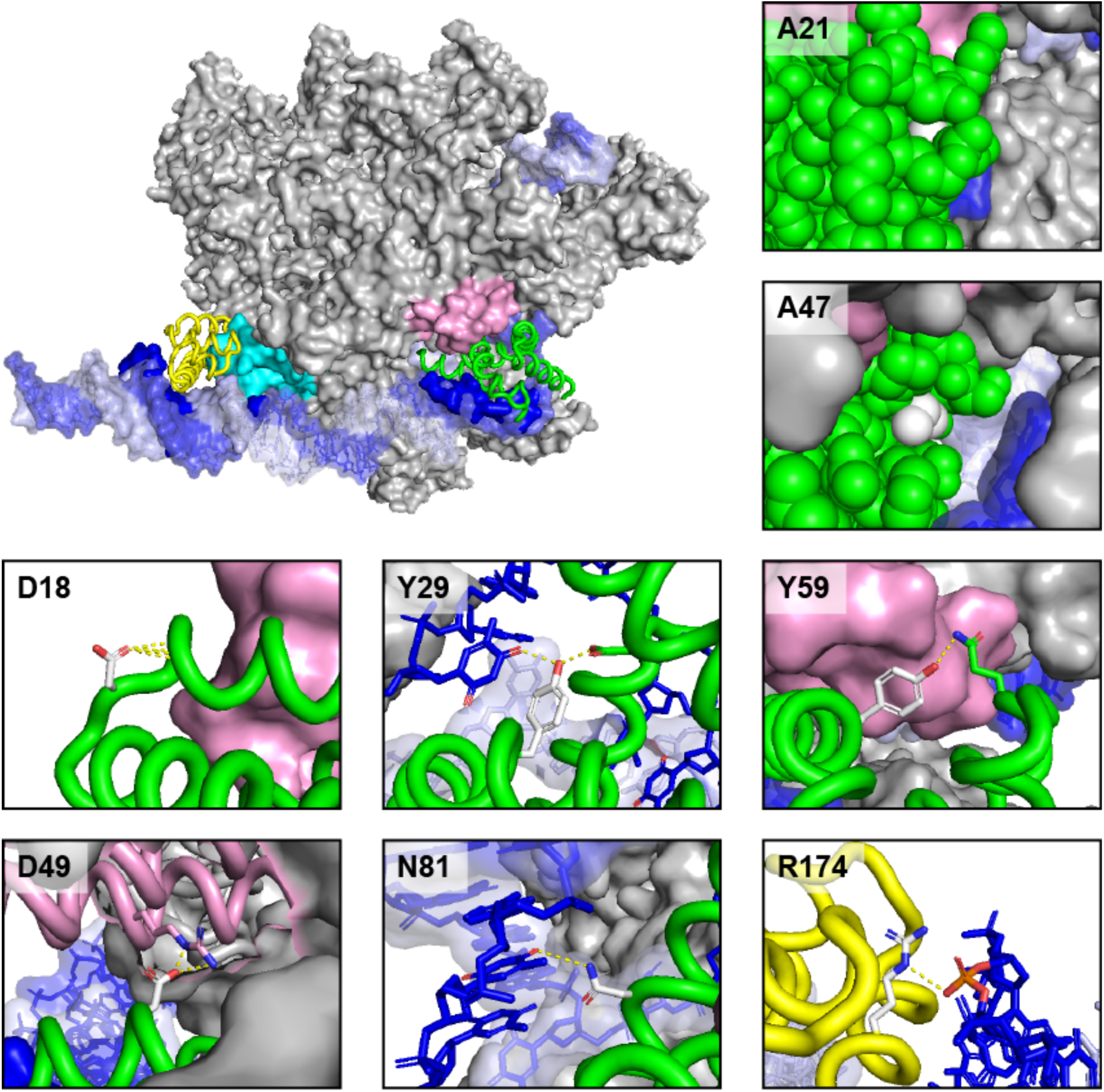
Location of affected AlgU residues in revertants that can grow in the absence of MucA. Model of the RNAP holoenzyme containing σ^E^ bound to the promoter. Gray, RNAP core; cyan, β flap; pink, β’ coiled coil; green, σ^E^ Region 2; yellow, σ^E^ Region 4; blue/light blue, promoter element. Insets, location of residues that are substituted in the revertants with missense *algU* mutations. The side chain of the affected residue is in white. Substitution of A21 and A47 (V in model) likely affect protein packing and folding of AlgU. Substitution of D18, Y29, and Y59 would reduce predicted intra- and inter-molecular hydrogen bonding and likely affect AlgU folding. Substitution of D49, N81, and R174 would likely affect the interaction of AlgU with the RNAP core or the promoter. Dashed yellow lines, hydrogen bonds; red atoms, oxygen; blue atoms, nitrogen; orange, phosphorus.

**Figure S5.**
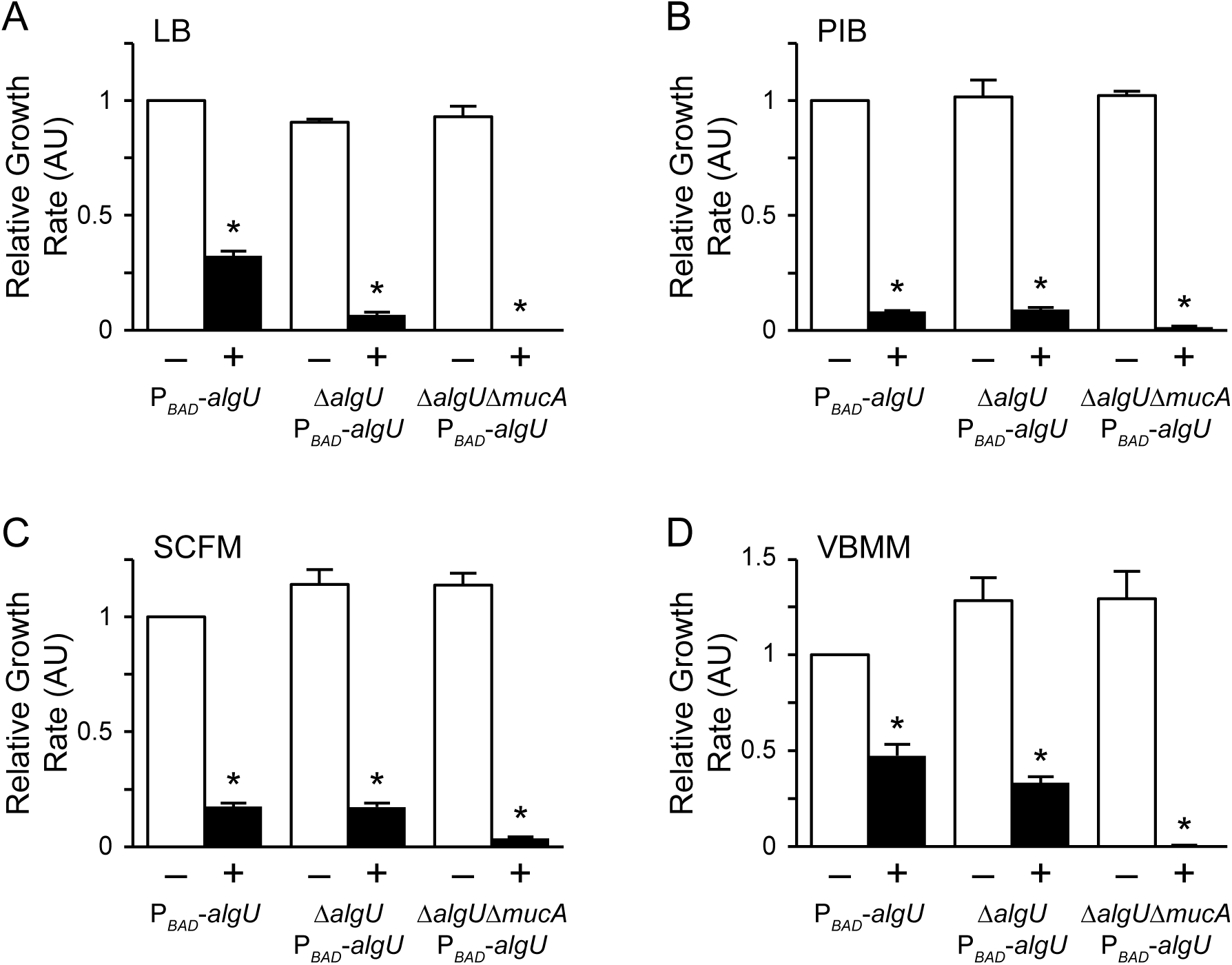
Overexpression of *algU* in the absence of *mucA* causes a growth defect in various media. Growth rate of indicated strains grown in (A) LB, (B) PIB, (C) SCFM, and (D) VBMM with (+) or without (–) 1% arabinose (relative to PAO1 *att*Tn7::P*_araBAD_*-*algU* grown in the absence of arabinose). Error bars, SEM (N=3). Asterisk, statistically different from the same strain grown in the absence of arabinose (p < 0.01, N = 3, two-way ANOVA with post-hoc Bonferroni). See Table S3 for full statistical comparisons. LB contains tryptone and yeast extract, both of which provide protein hydrolates, as the main carbon source. The main carbon source of PIB is also protein hydrolates, but from pancreatic gelatin digest. While PIB also contains glycerol, which can serve as a carbon source, it is not a preferred carbon source for *Pseudomonas aeruginosa*. SCFM is a defined medium that contains amino acids as the primary carbon source, and mimics the nutrients found in the CF lung environment. VBMM is a minimal medium that contains citrate as the sole carbon source.

**Figure S6.**
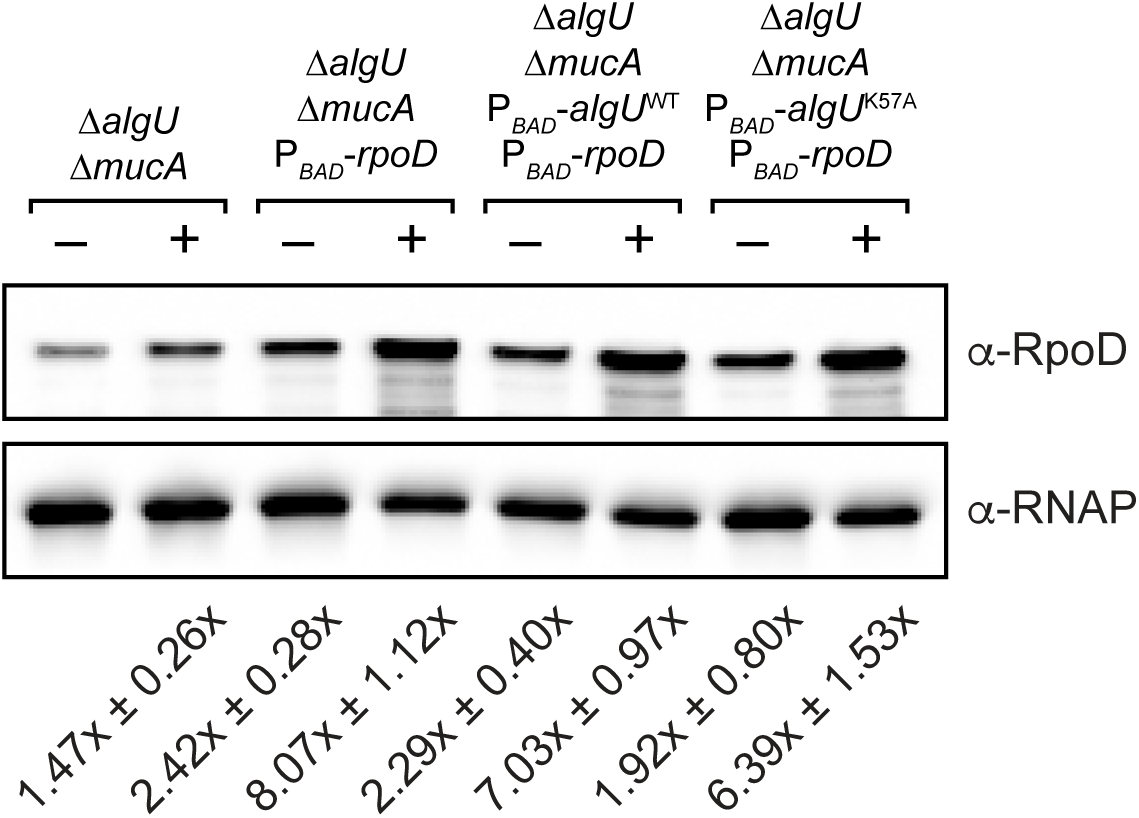
RpoD production in strains with an ectopic inducible *rpoD* allele. Expression of *rpoD* was inferred via semi-quantitative Western blot in the indicated strains (top). RNAP was used as a loading control (bottom). Strains were grown in LB with (+) or without (–) 2% arabinose. The relative amount of RpoD is indicated, where 1.00x represents RpoD levels in the parental strain without arabinose (± SD; N=3). While the strains produce significantly more RpoD in the presence of arabinose relative to the same strain in the absence of arabinose (p < 0.01), there is no statistical difference in the amount of RpoD produced from the induced strains containing an ectopic *rpoD* allele (two-way ANOVA with post-hoc Bonferroni).

**Table S1.** Eliminating alginate biosynthesis does not alleviate *mucA* essentiality.

| PAO1 | Number of isolates resolved to |  |
| --- | --- | --- |
| | WT | $\Delta mucA$ |
| WT | 168 | 0* |
| $\Delta algD$ | 144 | 0* |
| $\Delta algB$ | 158 | 0* |
| $\Delta algR$ | 153 | 0* |
| $\Delta amrZ$ | 131 | 0* |
| $\Delta algB \Delta algR \Delta amrZ$ | 133 | 0* |

\* p < 0.0001, Fischer's exact test

**Table S2.** Description of changes found in *algU* of PAO1 Δ*mucA att*Tn7::P*_rhaBAD_*-*mucA* revertants that can grow in the absence of *mucA* expression.

|  | Isolate | Genomic Change to *algU*<sup>†</sup> | Effect on AlgU<sup>‡</sup> |
| --- | --- | --- | --- |
| Multi-base pair deletions |  |  |  |
|  | #1-10 | 831242-831394 del | No product made |
|  | #2-9 | 831298-831491 del | No product made |
|  | #2-7 | 831467-831473 del | I56fs X97 |
|  | #2-11 | 831469-831502 del | K57fs X88 |
|  | #2-6 | 831595-831715 del | V99fs X107 |
| Single base pair deletions |  |  |  |
|  | #2-14 | 831467 del | I56fs X99 |
|  | #1-1 | 831510 del | Y72fs X99 |
|  | #2-10 | 831538 del | I80fs X99 |
| Nonsense mutations |  |  |  |
|  | #2-1 | 831433 C>T | Q45X |
|  | #2-8 | 831521 G>A | W74X |
| Multi-base pair insertions |  |  |  |
|  | #1-9 | 831427-831435 ins GACGCCCAG | D43-A44 ins DAQ |
|  | #1-8 | 831430-831438 ins GCCCAGGAA | A44-Q45 ins AQE |
|  | #1-11 | 831436-831444 ins GAAGCCCAG | E46-A47 ins EAQ |
|  | #2-4 | 831436-831444 ins GAAGCCCAG | E46-A47 ins EAQ |
|  | #1-5 | 831445-831456 ins GACGTAGCCCAG | D49-V50 ins DVAQ |
|  | #1-6 | 831451-831459 ins GCCCAGGAA | A51-Q52 ins AQE |
| Missense mutations |  |  |  |
|  | #1-3 | 831353 A>G | D18G |
|  | #2-12 | 831362 C>T | A21V |
|  | #2-3 | 831386 A>G | Y29C |
|  | #2-2 | 831439 G>A | A47T |
|  | #1-2 | 831446 A>G | D49G |
|  | #2-13 | 831446 A>G | D49G |
|  | #1-7 | 831476 A>G | Y59C |
|  | #2-5 | 831541 A>G | N81D |
|  | #1-4 | 831820 C>G | R174G |

<sup>†</sup> genomic location based on the Pseudomonas Genome Database (Winsor *et al.*, 2016) for PAO1; bp, base pair; del, deletion; ins, insertion.
<sup>‡</sup> fs, frameshift; ins, insertion; X, stop codon.

**Table S3.**
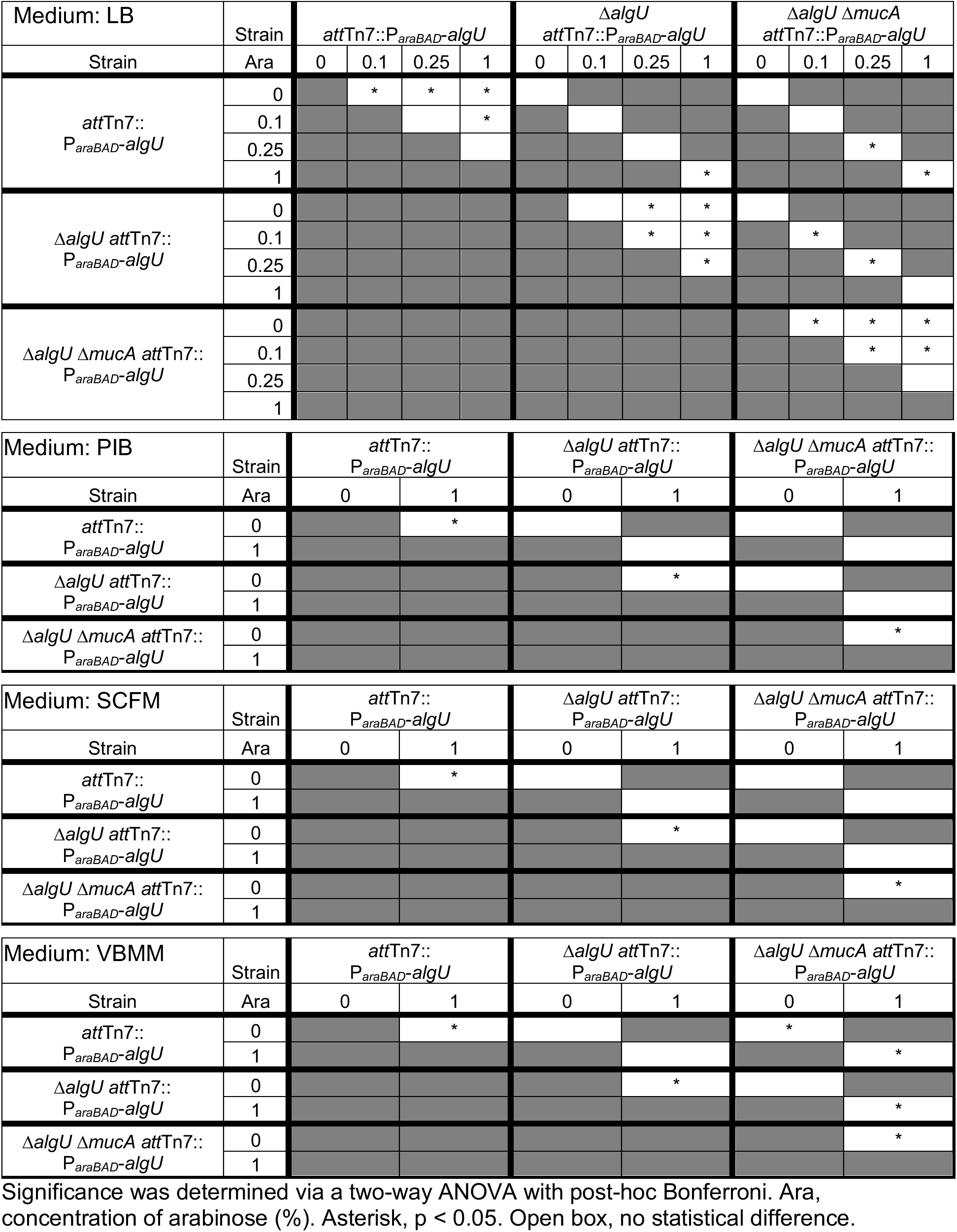
Statistical differences in growth rate among groups under conditions of *algU* induction.

**Table S4.** Statistical differences in growth rate among groups overexpressing *algU* and/or *rpoD*.

| | Strain | $\Delta algU \Delta mucA$<br>$P_{BAD-rpoD}$ | | $\Delta algU \Delta mucA$<br>$P_{BAD-algU^{WT}}$ | | $\Delta algU \Delta mucA$<br>$P_{BAD-algU^{WT}}$<br>$P_{BAD-rpoD}$ | | $\Delta algU \Delta mucA$<br>$P_{BAD-algU^{K57A}}$ | | $\Delta algU \Delta mucA$<br>$P_{BAD-algU^{K57A}}$<br>$P_{BAD-rpoD}$ | |
| --- | --- | --- | --- | --- | --- | --- | --- | --- | --- | --- | --- |
| Strain | Ara | 0 | 2 | 0 | 2 | 0 | 2 | 0 | 2 | 0 | 2 |
| $\Delta algU \Delta mucA$ | 0 | | | | | | | | | | |
|  | 2 |  |  |  | * |  |  |  | * |  |  |
| $\Delta algU \Delta mucA$<br>$P_{BAD-rpoD}$ | 0 | | | | | | | | | | |
|  | 2 |  |  |  | * |  |  |  | * |  |  |
| $\Delta algU \Delta mucA$<br>$P_{BAD-algU^{WT}}$ | 0 | | | | * | | | | | | |
|  | 2 |  |  |  |  |  | * |  | * |  | * |
| $\Delta algU \Delta mucA$<br>$P_{BAD-algU^{WT}}$<br>$P_{BAD-rpoD}$ | 0 | | | | | | * | | | | |
|  | 2 |  |  |  |  |  |  |  |  |  |  |
| $\Delta algU \Delta mucA$<br>$P_{BAD-algU^{K57A}}$ | 0 | | | | | | | | * | | |
|  | 2 |  |  |  |  |  |  |  |  |  |  |
| $\Delta algU \Delta mucA$<br>$P_{BAD-algU^{K57A}}$<br>$P_{BAD-rpoD}$ | 0 | | | | | | | | | | * |
|  | 2 |  |  |  |  |  |  |  |  |  |  |

Significance was determined via a two-way ANOVA with post-hoc Bonferroni. Ara, concentration of arabinose (%). Asterisk, $p < 0.05$ . Open box, no statistical difference.

